# The 3D architecture of the pepper (*Capsicum annum*) genome and its relationship to function and evolution

**DOI:** 10.1101/2021.12.10.470457

**Authors:** Yi Liao, Juntao Wang, Zhangsheng Zhu, Yuanlong Liu, Jinfeng Chen, Yongfeng Zhou, Feng Liu, Jianjun Lei, Brandon S. Gaut, Bihao Cao, J.J. Emerson, Changming Chen

## Abstract

The architecture of topologically associating domains (TADs) varies across plant genomes. Understanding the functional consequences of this diversity requires insights into the pattern, structure, and function of TADs. Here, we present a comprehensive investigation of the 3D genome organization of pepper (*Capsicum annuum*) and its association with gene expression and genomic variants. We report the first chromosome-scale long-read genome assembly of pepper and generate Hi-C contact maps for four tissues. The contact maps indicate that 3D structure varies somewhat across tissues, but generally the genome was segregated into subcompartments that were correlated with transcriptional state. In addition, chromosomes were almost continuously spanned by TADs, with the most prominent found in large genomic regions that were rich in retrotransposons. A substantial fraction of TAD boundaries were demarcated by chromatin loops, suggesting loop extrusion is a major mechanism for TAD formation; many of these loops were bordered by genes, especially in highly repetitive regions, resulting in gene clustering in three dimensional space. Integrated analysis of Hi-C profiles and transcriptomes showed that change in 3D chromatin structures (e.g. subcompartments, TADs, and loops) was not the primary mechanism contributing to differential gene expression between tissues, but chromatin structure does play a role in transcription stability. TAD boundaries were significantly enriched for breaks of synteny and depletion of sequence variation, suggesting that TADs constrain patterns of genome structural evolution in plants. Together, our work provides insights into principles of 3D genome folding in large plant genomes and its association with function and evolution.

## Introduction

The folding of chromosomes into higher-order chromatin domains ^1^, also known as topologically associating domains (TADs), appears to be conserved in evolution ^2^. TADs and similar structures occur in diverse groups of eukaryotes, from fungi and bacteria to plants and animals ^3^. It has been proposed that TADs are formed by varying mechanisms, of which loop extrusion and compartmentalization are two leading models in animal systems ^4–7^. While evidence suggests that these mechanisms may operate in tandem to jointly establish or maintain the spatial organization of the genome, the prevalence of each differs across species ^8–11^. Like animals, TAD-like domains have been observed in the Hi-C analyses of many plants; however, the mechanisms by which they form (and whether they are shared with animals) is largely unknown ^2,12^. Additionally, TADs organized by different mechanisms may exhibit distinct structural and functional properties ^8,13–15^. Thus, clarifying the underlying mechanisms of TADs is necessary for further elucidating their functional specialization.

Spatial genome organization is strongly associated with transcription. Numerous studies at the organismal ^16,17^, tissue^18^, and cell type ^19–21^ levels have established that rearrangement of 3D chromatin organization (i.e. higher-order chromatin structures, such as loops, TADs, and compartments) is associated with changes in gene expression. However, many studies suggest that TADs are not required for *cis* regulatory interactions that activate normal gene expression ^22–24^, suggesting that 3D chromatin domains may primarily act as an architectural framework to facilitate gene regulation ^25^. Although many recent attempts have been made to study these phenomena in plants ^26–29^, the relationship between 3D genome organization and the regulation of transcription in plant systems remains elusive.

TADs are thought to behave as functional and structural units of the genome in evolution ^5^. In metazoans, chromosomal rearrangement breakpoints rarely occur within chromatin domain bodies, implying that disruption of TAD integrity is unfavorable and subject to purifying selection ^18,30–33^. Indeed, long-range promoter-enhancer contacts that form loops are known to constrain large-scale genome evolution ^17^. On the other hand, three-dimensional chromatin structure can also affect patterns of both somatic mutation ^34^ and genomic variants across evolutionary timescales ^35^. Given that the spatial organization of the genome affects organismal function, an open question in plant biology is: how does natural selection affect the acquisition and fate of mutations – particularly, structural variants – that alter spatial organization? In plants, even though 3D genome organization is thought to play an important role in the polyploidization process ^29,36–38^, our understanding of the relationship between chromatin structure and structural variants remains incomplete.

Unlike in animals, where TADs can be readily detected genome-wide, small plant genomes like *Arabidopsis thaliana* and its close relative *Arabidopsis lyrata* carry few such domains ^39^. However, other plant species with relatively large genome sizes do exhibit more pronounced chromatin domain architectures ^40–43^. Comparisons between plant species imply that TAD prevalence in plants may be associated with genome size or other sequence properties, like the linear distribution of genes, regulatory elements, and transposable elements ^12,44,45^. Consequently, 3D genome organization appears to exhibit greater diversity in plants than it does in metazoans. This may also be true of the mechanisms that contribute to TAD formation. For example, TAD-like domains in maize and tomato are rarely demarcated by DNA loops; TADs instead largely coincide with compartments, suggesting compartmentalization is the leading mechanism for chromatin domain formation in these species ^41^. In contrast, recent studies in wheat have reported that a large proportion of chromatin domains are demarcated by chromatin loops, suggesting that loop extrusion is the more prevalent mechanism in this species. Many other features such as transcription factors are also found to be associated with the formation of plant chromatin domains ^14,40^. Thus, in plants, there appears to be variation not only in the prevalence of topological domains but also their mechanism of formation.

In this study, we aim to better understand the diversity of three-dimensional genome architecture across plants and the roles of chromatin features on genome function and evolution. To do this, we investigate principles of 3D genome folding in pepper (*Capsicum annuum*). We chose pepper because it is widely cultivated, because it is a member of the economically critical Solanaceae and because its 3D domain architecture exhibits clearly defined TADs that span most of the genome, comparable to *Drosophila* and mammals (Extended Data Fig. 1). We first generated a long-read chromosome-level reference genome assembly for an elite pepper inbred line ‘CA59’ and then constructed Hi-C maps for four tissues, including leaf, bud, pulp and placenta. Given these data, we characterize the general properties of major chromatin features (i.e. subcompartments, TADs, and loops) of 3D genome organization, their differences across tissues, and mechanisms or sequence features underlying TAD formation. Further integrating with genomic variation and transcription data, finally, we investigate the potential roles of 3D genome organization in genome structural evolution and gene expression.

## Results

### A chromosome-scale genome assembly of *Capsicum annuum*

We chose to sequence the pepper (*Capsicum annuum* L.) inbred line CA59 (Supplementary Fig. 1) for its agronomic characteristics, such as high yield, broad-spectrum disease resistance, and abiotic stress tolerance ^46^. We performed *de novo* assembly of the genome using ∼452 Gb Pacific Biosciences (PacBio) long-read sequence data (150× genomic coverage), ∼362 Gb (120×) short-read sequence data (150 bp paired-end, BGI genomics), and ∼415 Gb (135×) Hi-C data (150 bp paired-end, BGI genomics) (Supplementary Table S1 and Extended Data Fig. 2). Assembling of PacBio long reads alone produced a draft assembly that had 633 gapless contigs with a contig N50 of 41.3 Mb (Supplementary Table S2). Such high continuity is likely a consequence of low heterozygosity (0.23%) in our sample and the length of reads (subread N50 is 28,351 bp). The draft assembly was polished with short reads until reaching an estimated Phred quality score of Q52. Using Hi-C linkage information, 505 out of the 633 initial contigs were scaffolded into 12 pseudomolecules (scaffold N50 is 262 Mb) spanning 3.07 Gb sequences, leaving 128 unplaced contigs occupying only 11.66 Mb sequences (Supplementary Table S2). Our chromosome-scale assembly showed high collinearity with the previous Zunla-1 assembly ^47^ (Extended Data Fig. 3a) whose contigs were ordered and oriented via a high density genetic map, providing corroborating evidence for the accuracy of the Hi-C scaffolding result. The total genome size of the final assembly was similar to the estimated value (∼2.95Gb) based on the k-mers frequency analysis (Extended Data Fig. 2) and previous studies of pepper accessions ^47–49^.

Assembly quality evaluation with 1,440 Benchmarking Universal Single-Copy Ortholog (BUSCO) genes (Embryophyta odb9 dataset) showed that our chromosome-scale assembly recovers 95.8% of these genes, exceeding all previous *Capsicum* genome assemblies that were based on only Illumina sequencing (Supplementary Table S3). Moreover, *de novo* annotation of long terminal repeat retrotransposons (LTR-RTs) identified between 2,917 and 4,285 more full-length elements in our assembly than for previous assemblies (Supplementary Table S4), a likely consequence of higher continuity and completeness of our assembly in intergenic regions. Our assembly represents the first reference-quality genome assembly for pepper exceeding the EBP 6.C.Q40 standard.

Gene annotation was conducted combining evidence from PacBio full-length mRNA sequencing data (Iso-Seq) generated from five tissues (leaf, bud, pulp, placenta, and root), protein sequences previously annotated in closely related genomes, and *ab initio* prediction (Supplementary Table S5). A total of 46,160 protein-coding genes were predicted, which were enriched towards the ends of the chromosomes (Extended Data Fig. 3b), resembling observations in other large plant genomes. Preservation of synteny between genomes of pepper and three distantly related solanaceous species (tomato, eggplant, and potato) was thus common at chromosome ends (Extended Data Fig. 3c). We also annotated repeat content. Approximately 84.71% of the pepper genome was annotated as repetitive sequences, of which LTR-RTs alone make up 73.21% (Supplementary Table S6), including 59.89 Mb (1.95%) that represent 7,074 full-length elements (Extended Data Fig. 4). This result suggests that the vast majority of LTR-RTs in the pepper genome are fragmented. Amongst annotated LTR-RTs, 7 families were abundant, with 50 or more copies in the genome per family, representing ∼2,430 total insertions. Interestingly, most insertions in each of these 7 families had identical 5’ and 3’ long terminal repeats, indicating recent bursts of retroposition. Additional structural analysis of LTR-RT elements along the chromosomes suggests illegitimate recombination is the major process driving the rapid decay of LTR-RTs in the pepper genome (Supplementary Results and Extended Data Fig. 4).

### Hi-C interaction maps from four tissues

To interrogate the 3D genome architecture of *C. annuum*, we generated in situ Hi-C data from four tissues including leaf, bud, pulp, and placenta, each with two biological replicates. A total of 5.54 billion raw Hi-C read pairs (2 × 150bp) were produced, ranging from 557 to 788 million reads across samples, corresponding to raw sequencing coverages from 54x to 77x (Supplementary Table S7). We constructed Hi-C maps using both HiCExplorer ^11^ (Supplementary Table S8) and Juicer ^50^ (Supplementary Table S9). All Hi-C maps achieved a resolution around or higher than 10 kb (Supplementary Table S10), following previously described methods ^8^. Quality assessment using 3DChromatin_ReplicateQC toolkit ^51^ shows that our Hi-C data are of high quality as evidenced by QuASAR quality scores (0.039-0.061) ^52^ (Supplementary Table S11) and agreements between replicates (Supplementary Table S12). The reproducibility of Hi-C maps between biological replicates was also supported by the Pearson correlation analysis of their contact frequencies (Extended Data Fig. 5a).

We then compared Hi-C maps generated from different tissues. Inspection of the Hi-C maps revealed that the contact density was strongly concentrated along the main diagonals (**Fig. 1a** and Extended Data Fig. 5b), suggesting 3D proximity of pairs of loci is highly correlated with their linear genomic distance, as expected. As in other large plant genomes ^41^, we observed an X-shaped trans-interaction pattern, though we observe it only in certain tissues, like leaf and bud and not pulp and placenta (**Fig. 1a** and see juicer Hi-C maps in Extended Data Fig. 5b). The anti-diagonal pattern has been suggested as reflective of the chromosome “Rabl’’ conformation within the nucleus ^53,54^. This phenomenon reflects not only the presence of long-distance interactions, but also a correlation between the length of the span of such long-distance associations with distance from the centromere. We observed significantly more (Wilcoxon matched pairs signed rank test *P* ∼0.0005) long-range (>20 Mb) versus short-range contacts (<20 Mb) in leaf and bud compared to pulp and placenta (**Fig. 1b-d** and Extended Data Fig.5c-e). These observations may be the underlying cause of the hierarchical classification of the Hi-C maps, wherein leaf and bud clustered together while pulp and placenta cluster together (Extended Data Fig. 5a). These results demonstrate how global chromosome conformation inside nuclei might differ between cells from different plant tissues.

**Fig. 1:**
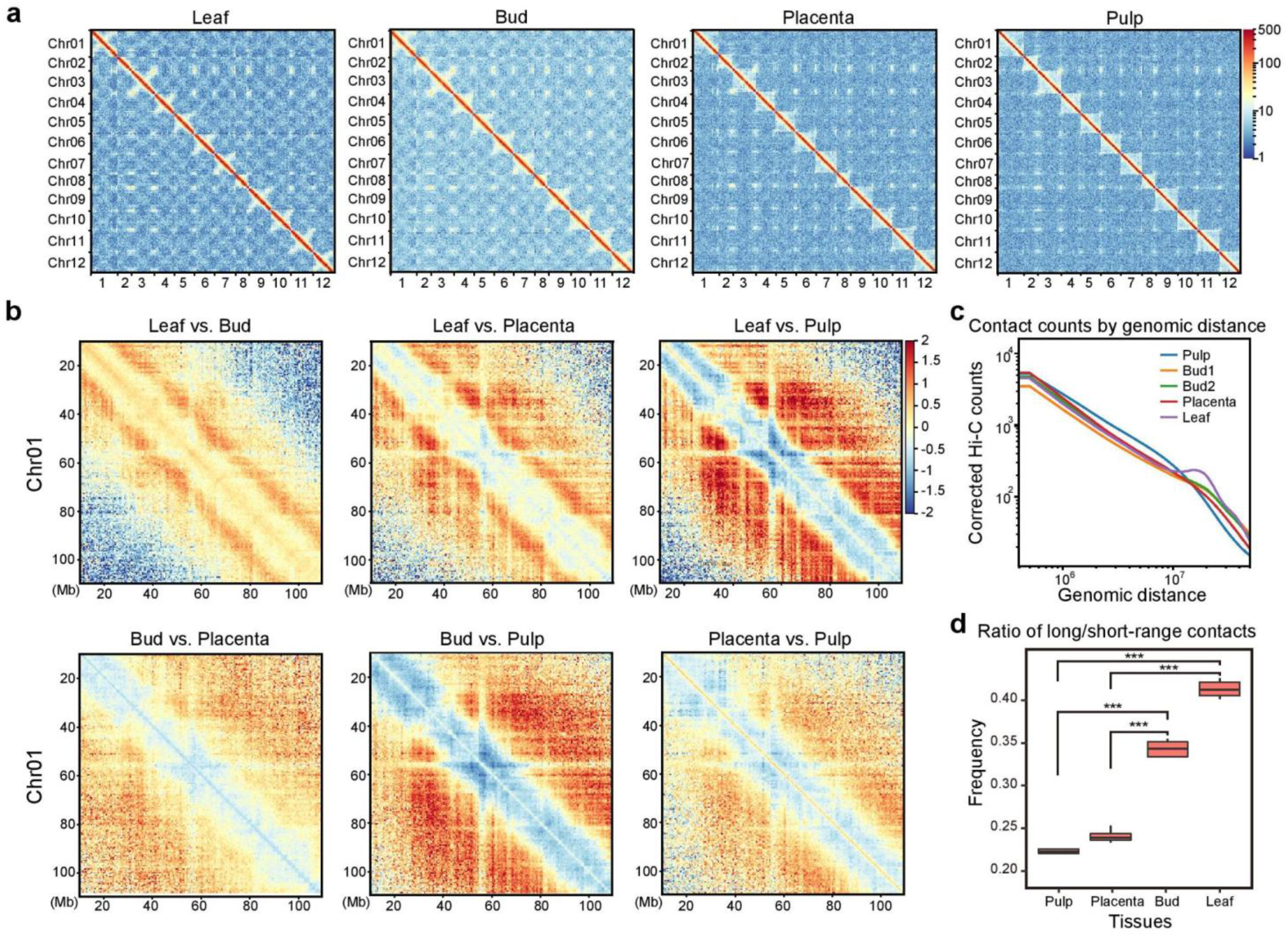
Hi-C interaction matrices (HiCExplorer) generated from four tissues of *C. annuum*. **a**, Genome-wide normalized and corrected Hi-C maps at 500kb resolution. In leaf and bud, an X-shaped trans-interaction signal appears, while it is weaker or not evident in contact maps of pulp and placenta. See also juicer Hi-C maps in Extended Data Fig. 5b. **b**, The log_2_-transformed ratio of Hi-C matrices between two tissues. Red designates enrichment in the first tissue, and blue depletion. **c**, The genomic distance *vs*. contact counts plot using Hi-C matrices at 500kb resolution. Leaf and Bud show an enrichment of long-range contacts (>20 Mb) than pulp and placenta. Only samples in the first batch were shown. See samples in the second batch in Extended Data Fig. 5c and results based on juicer Hi-C maps in Extended Fig. 5d. **d**, The ratio of long-range (>20 Mb) versus short-range contacts was calculated for each chromosome. This ratio is significantly higher in leaf and bud compared to pulp and placenta. *** indicates *p* < 0.0005, Wilcoxon matched pairs signed rank test. See also results obtained from juicer Hi-C maps in Extended Data Fig. 5e.

### Subcompartment patterning mirrors transcriptional state

At a global level, eukaryotic genomes are segregated into active (A) and inactive (B) compartments ^55^. Using an eigenvector analysis of the 500-kb Hi-C contact matrix after normalization by the observed/expected method ^1^, chromosomes were clearly divided into ‘A’ and ‘B’ compartments. ‘A’ compartments were generally distributed near telomere regions. ‘B’ compartments covered the large middle repetitive regions of chromosomes (**Fig. 2a**), consistent with the global distribution of gene density and repetitive sequence content. However, this method failed to resolve the identification of A/B compartments using Hi-C contact matrices at higher resolution (e.g. 40-kb).

**Fig. 2:**
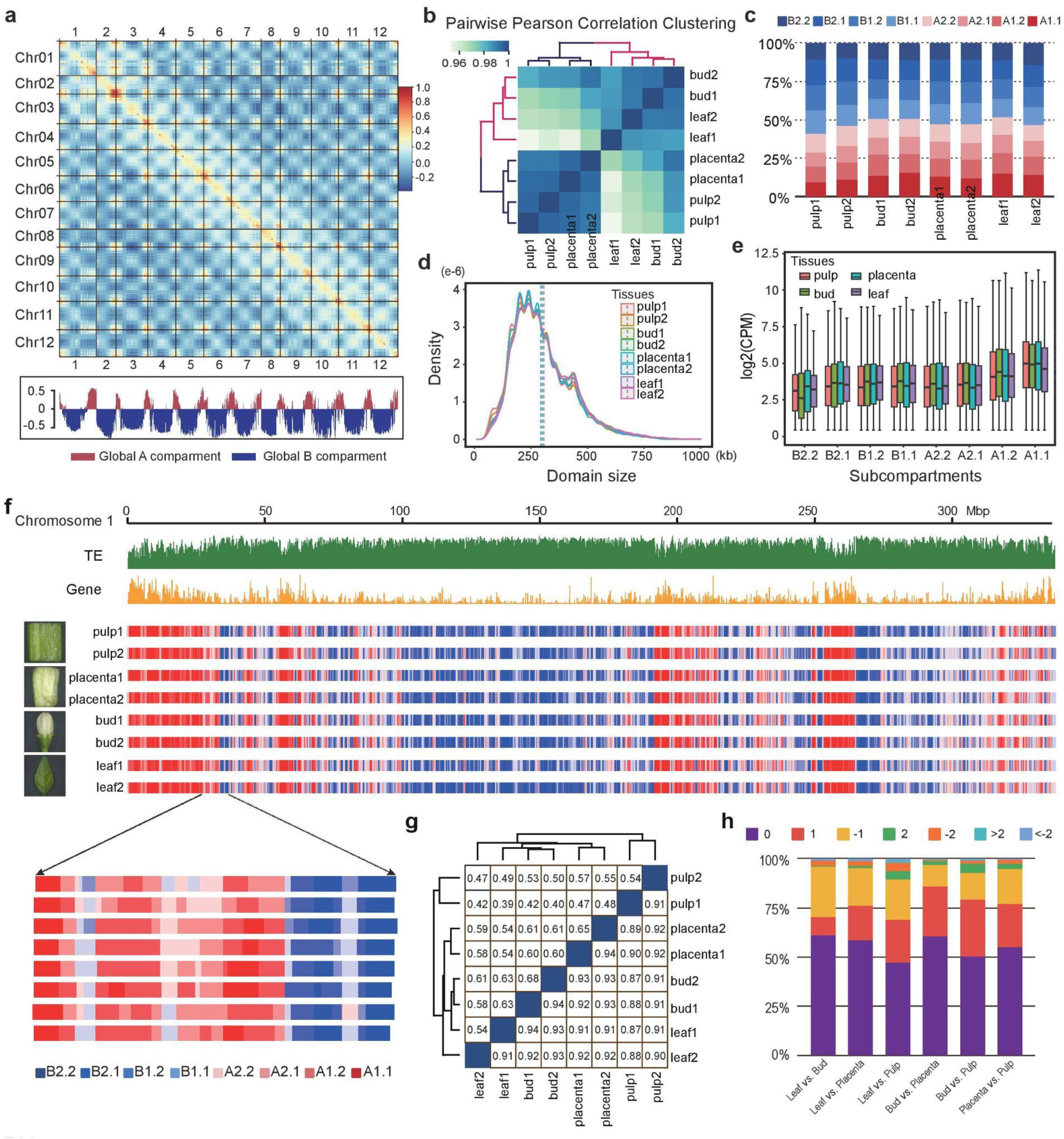
The inferred multi-scale subcompartments in the *C. annuum* genome are highly correlated with transcriptional landscape and maintained across tissues. **a**, Pearson correlation matrix heatmap at 500 kb resolution (leaf) shows segregation of the pepper genome into global A/B compartments. The first principal component (PC1) derived from analysis of this matrix was used to define the A and B compartments and is displayed below. Positive PC1 values are shown in red, representing A compartments, and negative PC1 values are shown in blue and designated as B compartments. **b**, Hierarchical clustering analysis of the Hi-C contact frequency (at 40 kb resolution) after batch effect correction across tissues and biological replicates. As expected, tissues are clustered together. **c**, The percentage of each Calder-inferred subcompartment (e.g. A1.1, A1.2, A2.1, A2.2, B1.1, B1.2, B2.1, and B2.2) across tissues. A and B compartments each occupy roughly half of the genome. **d**, The size distribution of the *Calder*-inferred subcompartments across tissues. All samples display a constant size distribution with a mean value of ∼300kb. **e**, The transcription levels, measured in counts per million (CPM), within subcompartments are generally consistent across tissues and correlate positively with subcompartment ranks from B2.2 to A1.1. **f**, Correspondence of subcompartments and the distribution of retrotransposon and gene density profiles. The example is shown for chromosome 1; for the other chromosomes see Extended Data Fig. 6c. **g**, Similarity of the A/B compartments and subcompartments between tissues. The upper part of the matrix is shown for subcompartments, and the lower part of the matrix is shown for A/B compartments. **h**, Subcompartment switching across four tissues. Pairwise comparisons across four tissues were shown. Numbers above where ‘0’ indicates unchanged subcompartment, ‘1’, ‘2’, and ‘>2’ indicate subcompartment shift spanning 1, 2 or more than 2 subcompartments for lower ranks to higher ranks, and ‘-1’, ‘-2’, and ‘<-2’ indicate indicate subcompartment shift spanning 1, 2 or more than 2 subcompartments for higher ranks to lower ranks.

To address this, we used a recent tool *Calder* ^13^ that allows us to infer compartments at high resolution (40-kb) at multiple scales. The Hi-C matrices were first normalized and corrected for unwanted variations, such as batch effects, using *BNBC* ^56^. The hierarchical clustering of the resulting Hi-C contact matrices showed high correlations within tissues (**Fig. 2b**). We ran *Calder* on the corrected Hi-C matrices with the default parameters that classify eight subcompartments. We found that roughly half of the genome is assigned as A subcompartments (i.e. A1.1, A1.2, A2.1, and A2.2), with the other half constituting B subcompartments (i.e. B1.1, B1.2, B2.1, and B2.2) (**Fig. 2c**). Across all subcompartments, decreasing rank can be assigned from A1.1, A1.2, …, B2.2. The average size of the inferred subcompartments is ∼300 kb, which is consistent for all samples (**Fig. 2d**).

We next investigated the biological properties of these multi-scale subcompartment assignments by analyzing their sequence features and transcription levels. We found that subcompartment rank is positively associated with gene content and negatively associated with retrotransposons content (Extended Data Fig. 6a). Consistent with this, subcompartment rank (from A1.1 to B2.2) is also positively associated with transcription level (**Fig. 2e** and Extended Data Fig. 6b). The genome-wide distribution pattern of genes and TEs, together with the inferred-compartments further confirmed these findings (**Fig. 2f** and see other chromosomes in Extended Data Fig. 6c). These results suggest that the pepper genome can be segregated into multi-scale subcompartments that are associated with the transcriptional landscape.

We also assessed the consistency of compartment assignments across tissues. At the A/B compartment level, we observed 87-94% of the genome shared the same compartment in comparisons between pairs of samples (**Fig. 2g**). For subcompartments, this rate dropped to between 39% and 65% (**Fig. 2g**). However, transitions that spanned more than 2 subcompartments (transitions from A1.1 to A1.2 had a scale of 1, and transitions from A1.1 to A2.1 had a scale of 2, and so on) accounted for less than 2.1% of the genome across all comparisons (**Fig. 2h**) suggesting subcompartments are often preserved across tissues.

### TADs are prominent in the pepper genome and maintained across tissues

Like in animals ^*57*^, TADs in the pepper genome are organized in a hierarchical fashion, such that small domains often reside within larger ones (**Fig. 3a**,**b**). This nested pattern of TAD poses challenges for their consistent identification ^58^. Indeed, we used the Hi-C data to annotate TADs based on three programs (Arrowhead, HiCExplorer, and TopDom) and found considerable variation in TAD calls (Extended Data Fig. 7a). For example, using a 40-kb resolution leaf Hi-C map, these three methods identified 1,780, 4,663, and 2,641 TADs, with medium sizes of 1,100 kb, 720 kb, and 1,067 kb, and occupying 56%, 98%, and 92% of the genome, respectively (Extended Data Fig. 7b,c). Even so, a substantial number of TADs (1,911) are consistently identified by at least two methods (Extended Data Fig. 7c), comparable to what we previously observed in *Drosophila* ^33^, and these TADs covered 55.4% of the genome. These results show that TADs is a prominent feature of genome architecture in pepper.

**Fig. 3:**
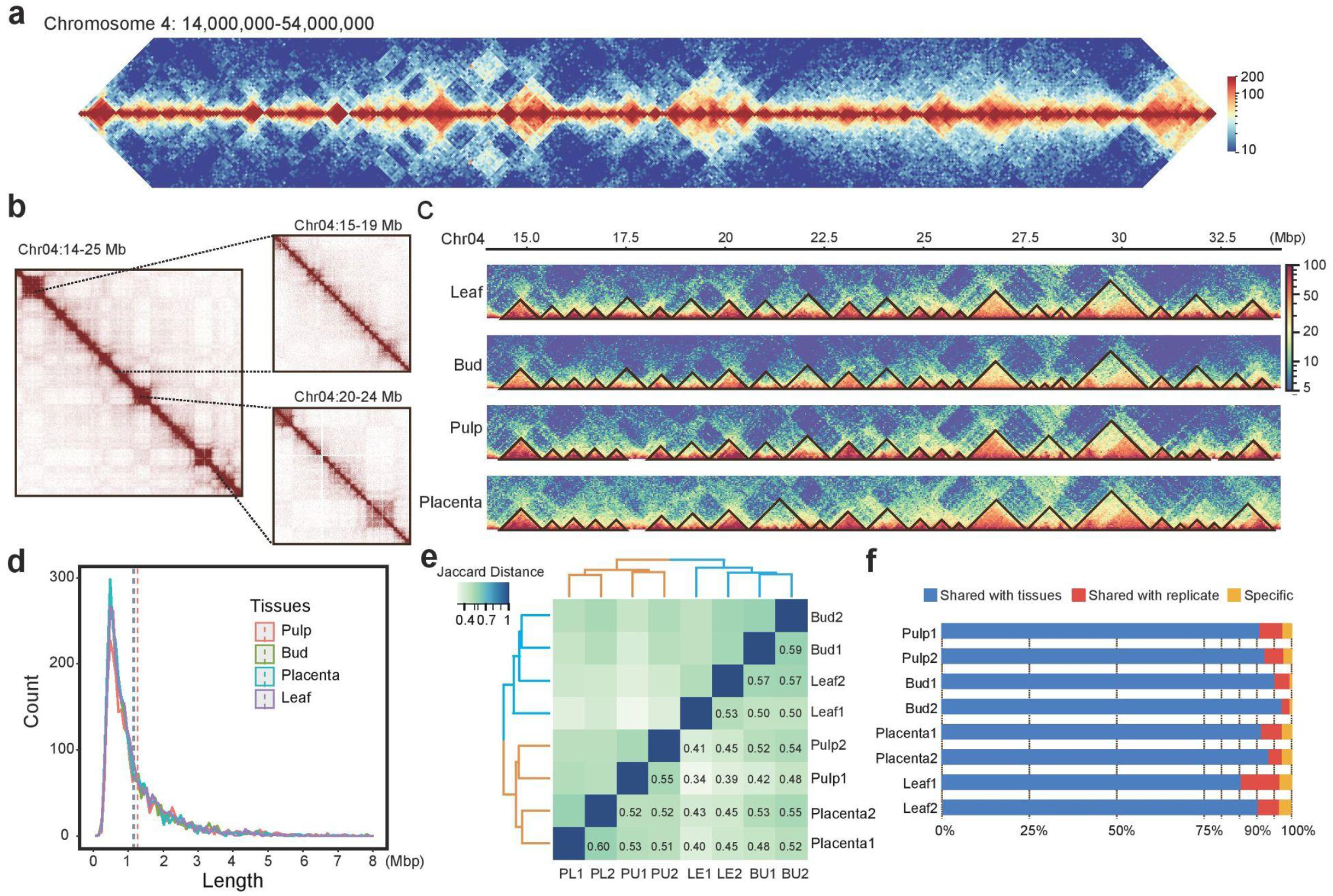
Prominent topologically associating domains (TADs) in the pepper genome. **a**, Pepper chromosomes are nearly completely segmented into TADs along their entire length. Example showing TAD structures (TADs are contiguous regions of enriched contact frequency that appear as squares in a Hi-C map) for a 40Mb region on chromosome 4. Leaf Hi-C interaction map at 100 kb resolution is shown. **b**, Small contiguous TADs show from Hi-C maps at higher resolutions, e.g. 40 kb (left), 10 kb (top right), and 5 kb (bottom right). **c**, TADs are highly conserved across the four studied tissues. TADs were annotated by TopDom using Hi-C maps at 40 kb resolution. **d**, Size distribution of the annotated TADs (TopDom). Vertical dashed lines indicate the mean values. **e**, Hierarchical clustering analysis of the called TADs based on the Jaccard distance of their shared genome coverage across tissues and biological replicates. As expected, tissues are generally clustered together. **f**, Conservation of TADs across tissues. Sample-specific TADs account for less than 5%. TADs annotated by TopDom at 40 kb resolution were used for this analysis.

By analyzing TADs annotated using TopDom, which performs well in TAD calling relative to most other methods ^58^, we found that our TAD calls were consistent across tissues in both profiles (**Fig. 3c**) and size (**Fig. 3d**). Hierarchical clustering analyses also demonstrated that TAD calls were reproducible across tissues and replicates (**Fig. 3e**). Roughly, between 58% and 79% of TADs (measured in their genome coverage), and between 60% and 91% of the boundaries were shared between any pairs of the samples (Extended Data Fig. 7d). At least 85% of TADs identified in one tissue were also detected in other tissues (**Fig. 3f**), suggesting TAD structures are highly conserved across the tissues investigated.

### Characterization and classification of TADs

We examined the relationship between TADs and compartment domains (CDs). In contrast to previous studies ^41^, we found that they rarely coincided (**Fig. 4a** and see more examples in Extended Data Fig. 8a), as shown by comparing TADs identified at multiple conditions to the *Calder*-inferred CDs (inferred from both 40-kb and 100-kb Hi-C matrices). In all comparisons, we found less than 11% of TADs coincide with CDs (i.e. exhibit ≥80% reciprocal overlap of segments, Extended Data Fig. 8b). This result suggests that compartmentalization may contribute to the formation of only a small fraction of TADs in the pepper genome ^6^.

**Fig. 4:**
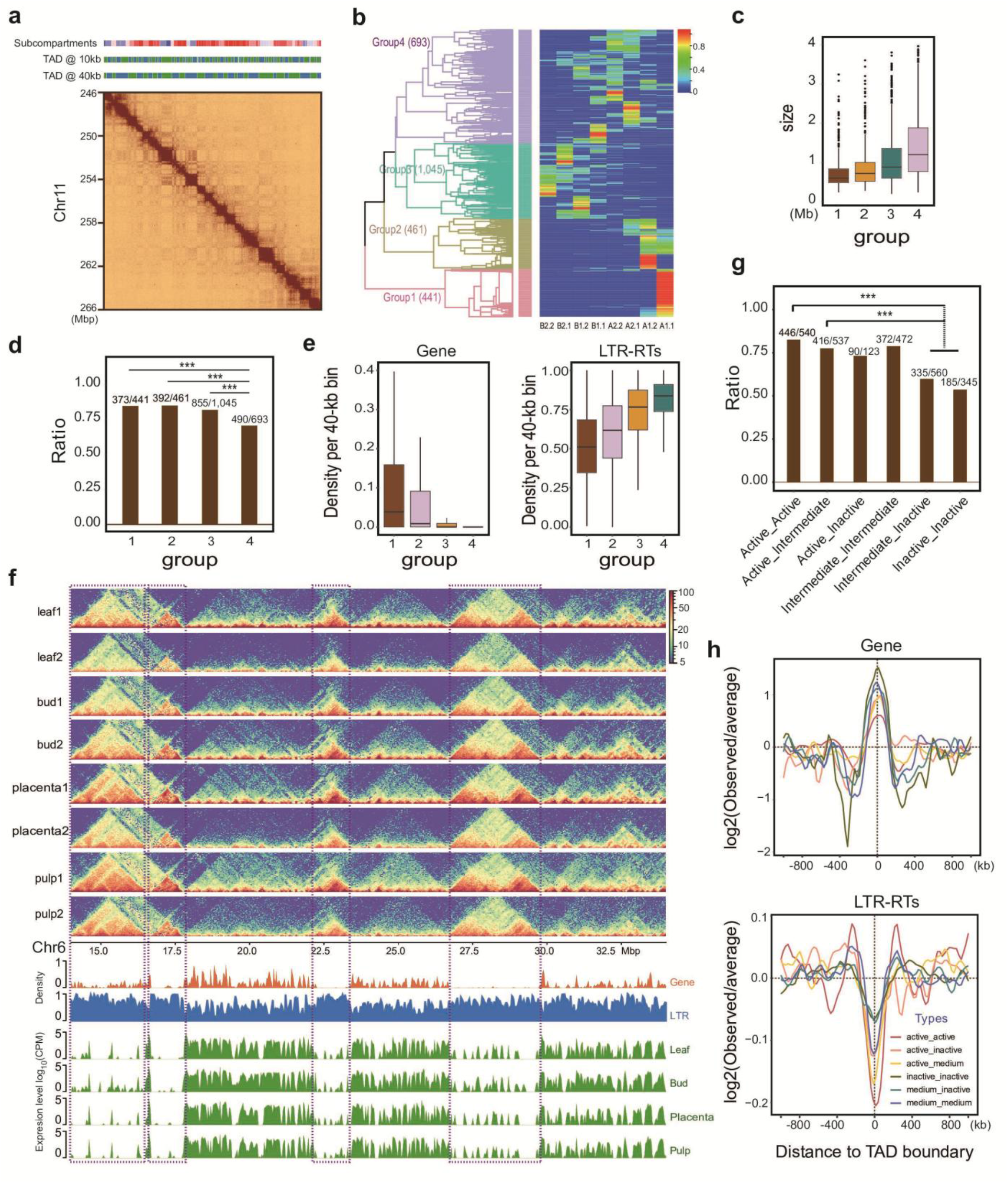
Characterization and categories of TADs in the pepper genome. **a**, TADs do not often coincide with compartments. Examples of comparison between *Calder*-inferred subcompartments (40-kb) and HiCExplorer-inferred TADs (10-kb and 40-kb) for a 20-Mb region on chromosome 11 (see panels above the Hi-C heatmap). Result was obtained from leaf Hi-C data. See also additional example regions in Extended Data Fig. 8a. **b**, Classification of TADs (for those annotated by TopDom using Hi-C map at 40-kb resolution) based on the enrichment of *Calder*-inferred subcompartments. See also the result for Arrowhead TADs in Extended Data Fig. 8c. **c**, Sizes vary across TAD groups. Generally, TADs enriched for B subcompartments are larger than A subcompartments. **d**, TADs enriched for A subcompartments (active group) are more conserved between tissues than those enriched for B subcompartments (inactive group). Conservation was defined as the percentage of TADs annotated in one tissus (here for leaf) also present in any other tissue. *** indicates *P*-value < 0.0001 based on Fisher’s exact Test. **e**, TADs in the active group have a higher gene density in their bodies than TADs in the inactive groups, while the trend is reverse for retrotransposons. **f**, Example of TAD appearance across all eight Hi-C maps (4 tissues) for a 20-Mb region on chromosome 6. Gene and retrotransposon density, along with transcription profiles from four tissues for this region were shown in below panels. The purple dashed rectangles show prominent TADs coincide with genomic regions enriched in retrotransposons. See also additional example regions in Extended Data Fig. 8d. **g**, Boundaries of TADs in the active group are more conserved between tissues than boundaries of TADs in the inactive group. Conservation was defined as the percentage of TAD boundaries annotated in one tissues (here for leaf) also present in any other tissue. *** indicates *P*-value < 0.0001 based on Fisher’s exact Test. **h**, TAD boundaries are enriched for genes and depleted of retrotransposons, which is consistent for all types of TAD boundaries based on their flanking TADs.

In metazoans, TADs can be classified into groups with distinct patterns of histone modifications and transcriptional activity ^57^. Given that *Calder*-inferred subcompartments are highly correlated with these properties ^13^, we next investigated whether they facilitate biologically meaningful classification of TADs. *Calder* assigns each locus (i.e. 40-kb bin) as belonging to one of the 8 subcompartments. Using this information and a hierarchical clustering approach, we partitioned TADs (derived from 40-kb bins) into 4 clearly separated groups (Fig. 4b and Extended Data Fig. 8c). TADs in each group were significantly enriched for 1 or 3 subcompartments. In group 1 (n=441) and group 2 (n=461), TADs were composed of loci that were significantly enriched for A1.1 and A1.2 subcompartments, respectively. In group 3 (n=1,045), TADs were enriched for B1.1, A2.2 and A2,1 subcompartments. Finally, TADs in group 4 (n=693) were enriched for B2.2, B2.1, and B1.2 subcompartments. Although TADs do not always coincide with subcompartments, these results suggest that TADs tend to contain genomic bins of somewhat uniform subcompartments and with similar transcription levels.

To reflect the relationship between subcompartments and transcription levels, we labelled TADs in group 1 and group 2 as active TADs, TADs in group 3 as intermediate, and TADs in group 4 as inactive. We found that active TADs were smaller (medium length of 734 kb) than both intermediate (medium length of 1.01 Mb) and inactive (medium length of 1.3 Mb) TADs (**Fig. 4c**), and active TADs tended to be more conserved across tissues than inactive TADs (**Fig. 4d**). Additionally, these observations were also consistent with their association with gene and LTR-RT content (**Fig. 4e**). Notably, the most prominent TADs (i.e. clearly visible as large squares in the Hi-C maps) formed in genomic intervals with a long stretch of enriched LTR-RTs and were always flanked by active transcription regions (**Fig. 4f** and Extended Data Fig. 8d). This pattern supports the hypothesis that the existence or formation of TAD structure in plants is associated with certain sequence features, such as genes, regulatory elements, and transposable elements ^12,44,45^.

The TAD boundaries of active TADs tended to be more stable across tissues than boundaries of inactive TADs (**Fig. 4g**), consistent with previous findings that active TADs are more conserved evolutionary across *Drosophila* species ^33^, further implying that active TADs are enriched for function relative to inactive ones. Consistent with our finding that TADs were typically flanked by regions of active transcription, we found that TAD boundaries to be enriched for genes and depleted for LTR-RTs (Extended Data Fig. 8d) and such a trend holds for all types of TAD boundaries (**Fig. 4h**).

### TAD boundaries are often demarcated by chromatin loops

To infer the potential mechanisms underlying TAD formation in pepper, we next attempted to annotate chromatin loops in the four studied tissues with combined Hi-C data from replicates. Using hicDetectLoops ^11^, we identified 5,746, 5,990, 7,701, and 9,142 chromatin loops in pulp, leaf, bud, and placenta, respectively, by merging output derived from Hi-C maps at multiple resolutions (e.g. 10 kb, 15 kb, 20 kb, and 25 kb) (Supplementary Table S13). Increased resolutions often resulted in larger loops but the vast majority (∼86%) of loops identified were <2 Mb apart (**Fig. 5a**), which is similar to humans ^8^. Approximately half of loops identified in one tissue were detected in other tissues (**Fig. 5b**). Combining loops identified from all four tissues resulted in a non-redundant set containing 19,521 loops. Among them, 5,728 were shared at least in two tissues and 13,793 were unique to a specific tissue. Importantly, when loops detected in one tissue were missing in another, we could not exclude the possibility that they were present but below the threshold of detection (Extended Data Fig. 9a). We reasoned that this might be due to technical limitations in loop detection approaches or reflect subtle changes in the interaction frequency between tissues ^16,20^. Therefore, we also employed Mustache, which also recovers loops with high levels of confidence ^59^. With Mustache, we identified 8,236 non-redundant loops, of which 5,282 (64.1%) were present in the hicDetectLoops calls. The set of 5,282 shared loops represent a conservative set supported by both annotation methods.

**Fig. 5:**
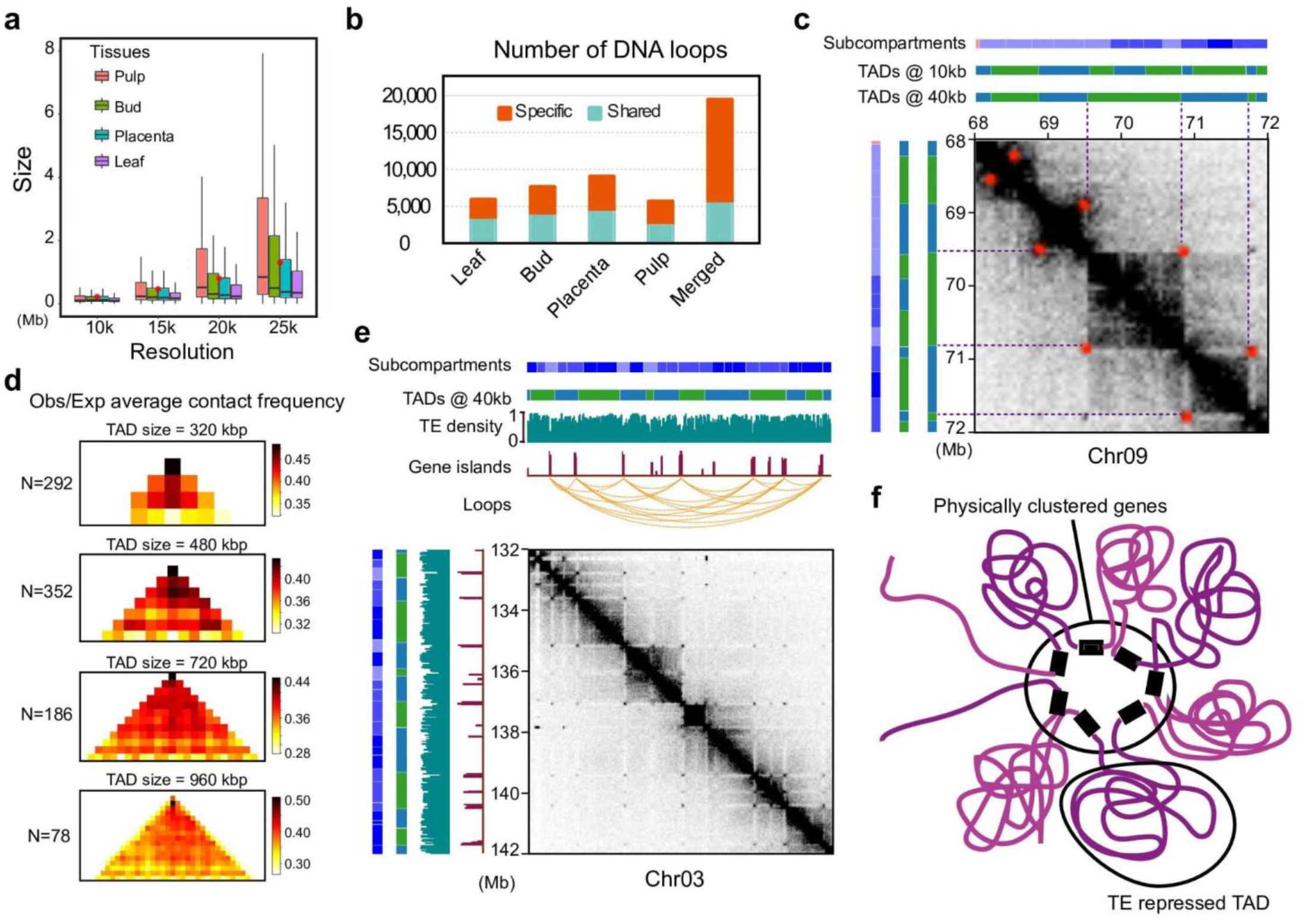
Chromatin loops frequently demarcate TADs and anchors of loops often coincide with genes. **a**, Chromatin loops identified across tissues with multiple resolutions (e.g. 10 kb, 15 kb, 20kb, and 25kb) by hicDetectLoops. Red dots indicate the mean size of loops. **b**, Numbers of tissue-specific and shared DNA loops. For each tissue, a shared loop was identified if it was present in any other tissues. A merged loop set was constructed by removing the redundant calls across all four tissues. **c**, Example showing a genomic region (Chr09: 68,000,000 - 72,000,000) where chromatin loops demarcate TADs. Subcompartments and TADs identified at both 10-kb and 40-kb resolution for this region were shown above and right. Loops were shown as red dots in the Hi-C contact maps at 40 kb resolution (leaf). **d**, Enrichment of contact frequency was observed at the corners of TADs of different sizes; that is, the peak loci are located at domain boundaries. More examples can be found in Extended Data Fig. 9d. **e**, Example showing loop anchors overlapping with gene islands within the repetitive genomic regions. More examples can be found in Extended Data Fig. 9e. **f**, Schematic representation of hypothesized gene-to-gene chromatin loops that mediate the formation of physically clustered genes.

In humans, chromatin loops frequently demarcate TADs—that is, the two anchors of a loop coincide with the two boundaries of a TAD ^8^. Based on the set of 8,236 loops called by Mustache, we found that this pattern was also very common in the pepper genome (**Fig. 5c** and Extended Data Fig. 9b,c). We found that a large fraction (31.4%) of loop anchors (Mustache calls) coincided with TAD boundaries (HiCexplorer TADs identified at 10 kb resolution Hi-C map of leaf), compared to 5% by random chance (*P*-value < 2.2 × 10^−16^, Fisher’s exact test). Similarly, ∼23% of TADs had loop anchors in their boundaries, compared to 3.9% by random chance (*P*-value < 2.2 × 10^−16^, Fisher’s exact test). This result contrasts with results from *Drosophila*^*10*^ and other plant species^10,41^ by suggesting that loop extrusion may act as a major mechanism that drives the formation of TADs in pepper. This suggestion is further supported by the elevated contact frequency between the two edges of a TAD, which shows up as high density in the anti-diagonal corners of TADs (**Fig. 5d** and Extended Data Fig. 9d).

We also observed that gene bodies are enriched at loop anchors (∼60% of loop anchors overlap with genes, compared to 29.4% at random, *P*-value < 2.2 × 10^−16^, Fisher’s exact test). In some cases, loops formed connecting gene islands that were several megabases apart in the highly repetitive regions (**Fig. 5e** and Extended Data Fig. 9e). Moreover, anchors of these loops often reciprocally overlapped with each other, indicating they collocated at a single spatial position, similar to loops in humans ^8^. We propose that such a configuration may facilitate the segregation of a physical cluster of genes into the active A compartments (**Fig. 5f**). These genes may share a chromatin-level co-regulation system and similar function ^60^.

### Breaks of synteny preferentially occur near TAD boundaries, despite high evolutionary conservation

TAD boundaries were enriched for sequence conservation at the nucleotide level, consistent with their enrichment for genes. For example, we found that DNA sequences at TAD boundaries exhibited roughly onefold enrichment of conservation relative to the genome-wide average when aligning conserved syntenic sequences from potato, tomato, and eggplant genome to the pepper genome (**Fig. 6a**,**b** and Extended Data Fig. 8e).

**Fig. 6:**
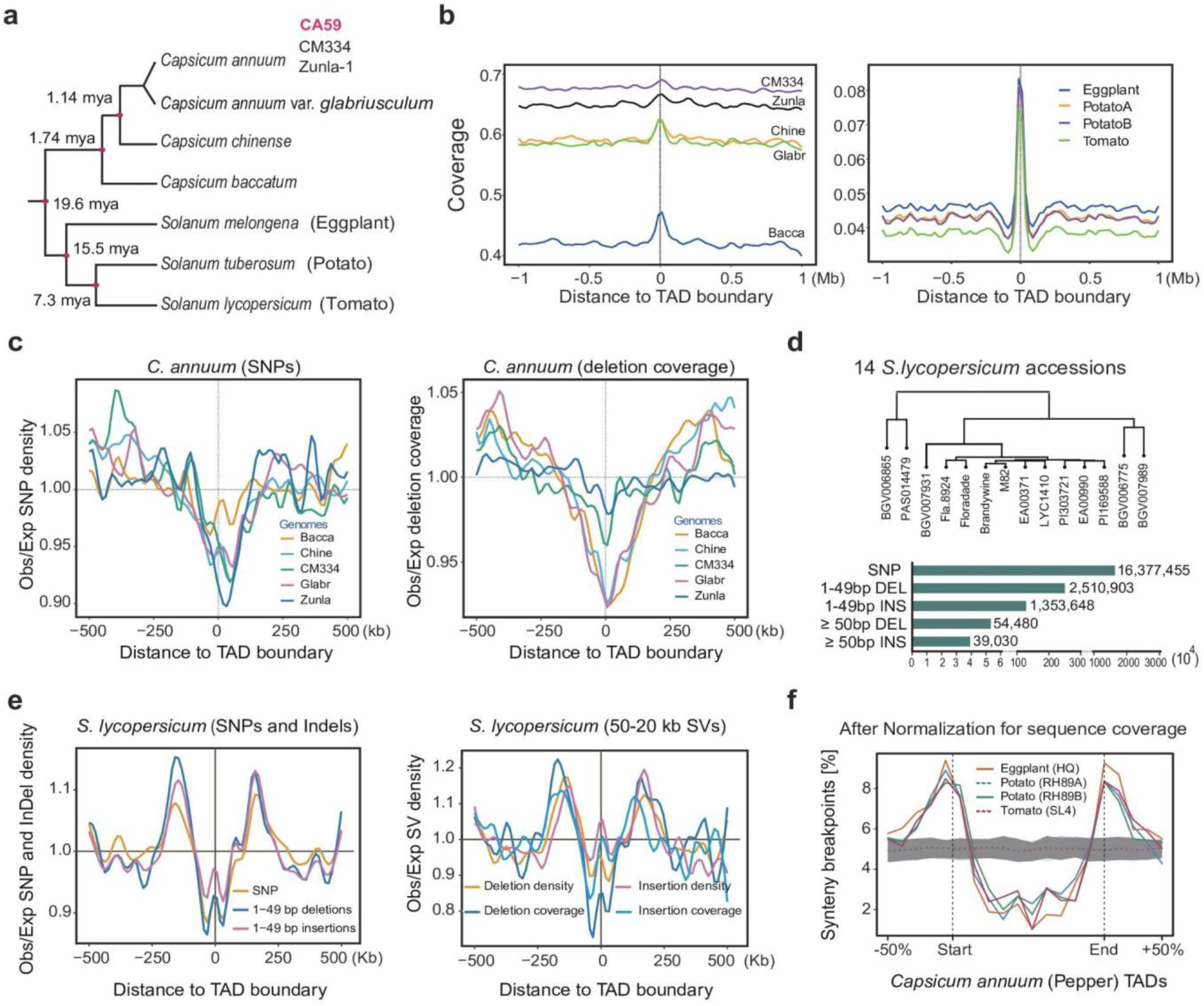
Pattern of genomic structural variants around TAD boundaries. **a**, Phylogenetic relationship of the studied *Capsicum* species and distantly related solanaceous species (*S. melongena, S. tuberosum*, and *S. lycopersicum*). The estimated divergence times were taken from ^66^ and ^48^. **b**, Alignable fraction (coverage) of syntenic and conserved genomic sequence around pepper TAD boundaries (TopDom calls). Left is shown for comparisons between CA59 and five closely related genomes, including two *C. annuum* accessions (*CM334* and *Zunla-1*), the wild progenitor of *C. annuum* (*C. annuum* var. *glabriusculum*), and two closely related species (*C. chinense* and *C. baccatum*). Right is shown for comparisons between CA59 and the three more distantly related solanaceous species. **c**, The observed (Obs) distribution of SNPs and deletions (coverage) near TAD borders relative to the expectation (Exp), based on the genomic background. SNPs and deletions were identified between CA59 and five closely related genomes. **d**, Genomic variants identified from high-continuous genome assembly of 14 *S. lycopersicum* accessions relative to the reference genome SL4 ^63^. **e**, The observed (Obs) distribution of SNPs, InDels, and large SVs (>50bp) near tomato TAD borders relative to the expectation (Exp), based on the genomic background. TADs were annotated by HiCExplorer with Hi-C data obtained from ^41^ using SL4 as the reference. **f**, TAD boundaries of pepper are enriched for evolutionary synteny breaks identified from distantly related solanaceous species. Dotted lines in gray show randomly simulated synteny breaks (n=100).

Given the above observation, we hypothesized that structural variations may also be constrained at such regions in plants, as observed in animals ^33,35^. To test this, we identified single-nucleotide variants (SNVs) and deletions relative to our CA59 assembly from five previous *Capsicum* assemblies, including two cultivated *C. annuum* accessions (CM334 and *Zunla-1*) and a wild progenitor (*C. annuum* var. *glabriusculum*), as well as two closely related species (*C. chinense* and *C. baccatum*) (**Fig. 6a**). With these data, we investigated the pattern of genome variation in the context of TADs and found that SNVs and deletions were strongly depleted around the pepper TAD boundaries (**Fig. 6c**). Such a pattern holds for TAD boundaries identified from all methods (Extended Data Fig. 10a,b). Since this pattern was not observed in rice TADs when assessing with population genomic variants ^61^, we asked whether it holds in more closely related species to *C. annuum*. We analyzed *S. lycopersicum* (tomato) using published Hi-C data and genomic variant calls identified from high-quality genome assemblies of 14 *S. lycopersicum* accessions ^62^ (**Fig. 6d**). Our analysis corroborates what we observed in pepper (**Fig. 6e**), suggesting constraints of genomic variations around TAD boundaries may be common across Solanaceae species.

In metazoans, TADs constrain large-scale genome evolution as indicated by the observation that breaks of chromosome rearrangements preferentially occur at TAD boundaries and are depleted in TAD bodies ^18,31,33^. Such a pattern, to our knowledge, has not yet been reported in plants. To examine this question, we identified genome synteny breaks between *C. annuum* and three distantly related Solanaceae species including, *S. lycopersicum* ^63^, *S. tuberosum* ^64^, and *S. melongena* ^65^ (**Fig. 6a**), which diverged from a common ancestor with *C. annuum* ∼19.6 million years ago. We found that synteny breaks were indeed enriched at TAD boundaries identified for each comparison between *C. annuum* and three solanaceous species (Extended Data Fig. 10c). This pattern persisted after normalization for sequence conservation level across TAD bodies (**Fig. 6f**). We also repeated the analyses using *S. lycopersicum* and *S. tuberosum* as a reference and obtained similar results, albeit with a weaker trend (Extended Data Fig. 10d,e). These results suggest that breaks of chromosomal rearrangements are enriched at TAD boundaries, despite high evolutionary conservation at such regions, in the Solanaceae.

### Changes in subcompartments, TADs, and loops are not associated with differential gene expression between tissues

To shed light on the relationship between 3D chromatin organization and gene expression, we tracked subcompartment switching to assess whether it corresponds to changes in the level of transcription and, if so, to what extent. To do so, we performed a pairwise comparison of both the subcompartment profiles and the transcriptomes of the four pepper tissues. In each pair comparison (for which there were six in total), all 40-kb bins (76,641, 3.07 Gb/40 kb, Supplementary Table S14) were assigned into three groups based on change status of subcompartments: (1) the ‘down’ bins in which subcompartment rank decreased by at least 1 (e.g. from A1.1 to A1.2 or lower), (2) the ‘up’ bins in which subcompartment rank increases by at least 1 (e.g. from B2.2 to B2.1 or higher), and (3) the ‘stable’ bins in which subcompartment remains unchanged. Correspondingly, we identified between 6,974 and 17,576 differentially expressed genes (DEGs, adjusted *P-value* < 0.01, among 38, 974 testable genes (with CMP > 0.05), Supplementary Table S14) in pairs of tissues using the R package Limma ^67^.

If changes in subcompartment patterning correlate with changes in gene expression, we expect that genomic regions with subcompartment switching contain more DEGs. Unexpectedly, we observed no enrichment of DEGs in either ‘up’ or ‘down’ bins compared to ‘stable’ bins in all six pairwise comparisons (Supplementary Table S14). However the percentage of genes with increased expression did rise from ‘down’ to ‘up’ bins, and the trend was reversed for genes that shifted from ‘up’ to ‘down’ bins (Supplementary Table S14). This trend was more pronounced when we only considered DEGs (Supplementary Table S14). Consistent with this, we observed that ‘up’ bins overlap genes that exhibited significantly higher log2(fold change) of transcriptional level than the ‘stable’ bins (Wilcoxon rank-sum test *p* < 0.006 for five comparisons; **Fig. 7a**), suggesting the ‘up’ bins are associated with increases in gene expression. In contrast, the ‘down’ bins overlapped genes that exhibit significantly lower log2(fold change) value than the ‘stable’ bins (Wilcoxon rank-sum test *p* < 0.023 for four comparisons; **Fig. 7a**), suggesting they are associated with decreases in gene expression. We repeated the analysis using transcription levels measured in bins (i.e. 40 kb) and obtained similar results (Supplementary Table S15 and Extended Data Fig. 11a). These results suggest that subcompartment patterning has limited effects on differential gene expression and may instead shape subtle changes in the amplitude of global transcription levels, especially for DEGs.

**Fig. 7:**
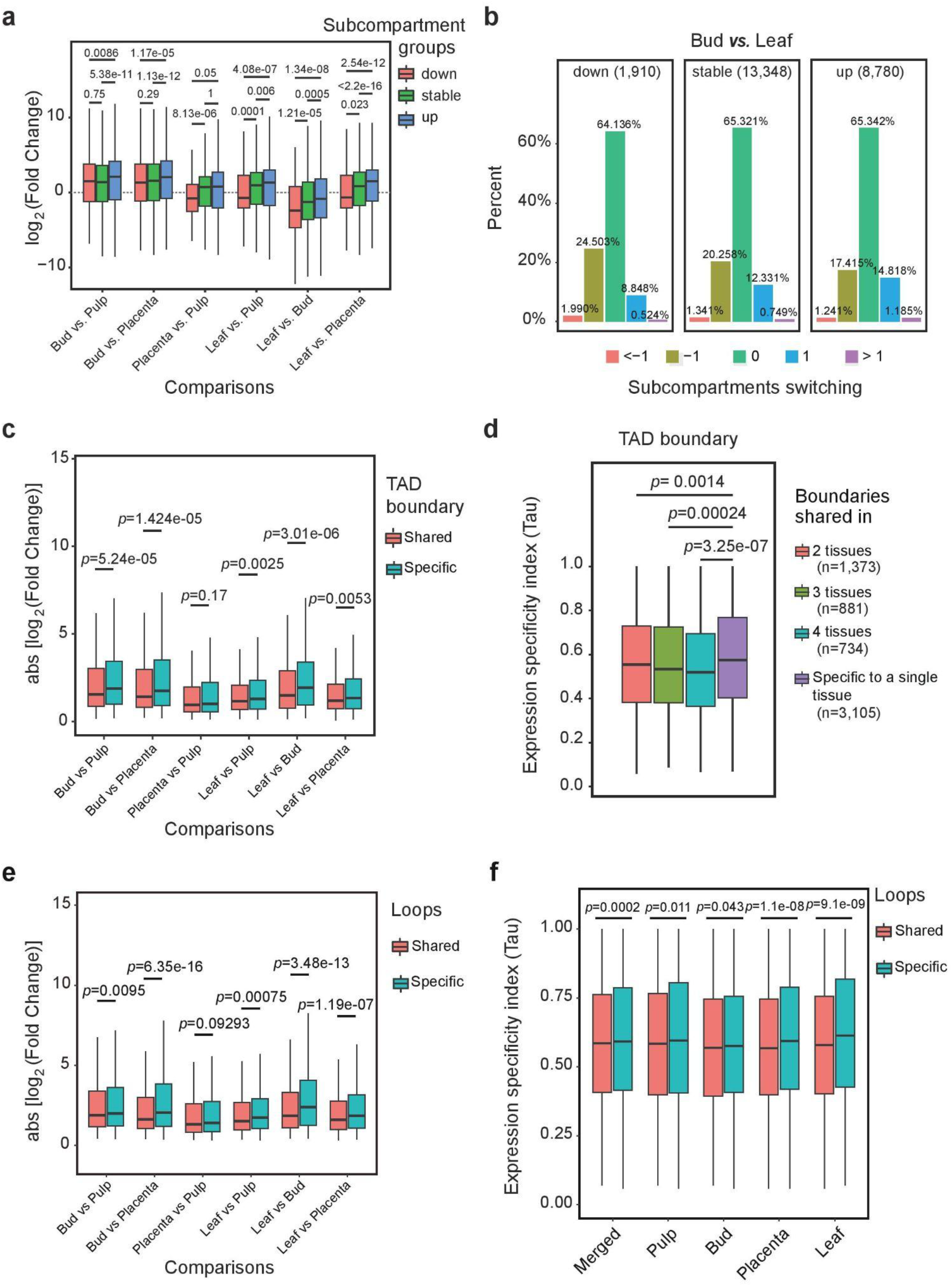
Subcompartments, TADs, and chromatin loops play roles in transcription stability. **a**, Genomic regions (i.e. 40-kb bins) switching from A to B compartments or from higher subcompartments to lower subcompartments (e.g. from A1.1 to A1.2) show a trend of decreasing expression, and conversely, switching from B to A compartment or from lower subcompartments to higher subcompartments show a trend of increasing expression. Pairwise comparisons of subcompartment shifts for expression profiles across leaf, bud, pulp and placenta are shown. **b**, 40-kb bins with decreased expression were slightly enriched for cases of subcompartment switching from a higher rank to lower ranks, while those with increased expression were slightly enriched for cases of subcompartment switching from lower ranks to higher ranks. Expression level decreases of more than 2 fold are labeled ‘down’, increases of more than 2 fold are labeled ‘up’, and changes within 2 fold are ‘stable’. Subcompartment switching from lower ranks to higher ranks are labeled ‘1’ if spanning 1 rank or ‘>1’ if more than 1 rank, from a higher rank to lower ranks are labeled ‘-1’ if spanning 1 rank or ‘<-1’ if more than 1 rank, and unchanged are labeled ‘0’. Specific comparisons can be found in Extended Data Fig. 11b. **c**, 40-kb bins overlapping with TAD boundaries that were conserved between tissues exhibit a relatively smaller absolute change fold in expression level than those overlapping with tissue specific TAD boundaries. Pairwise comparison across four tissues was conducted. Result depicted is for Arrowhead TAD annotation at 40 kb resolution, see results for other methods in Extended Data Fig. 12e. **d**, 40-kb bins overlapping with shared TAD boundaries across tissues exhibit a significantly lower expression specificity index Tau value compared to those overlapped with tissue-specific TAD boundaries. **e**, 40-kb bins overlapping anchors of shared chromatin loops between tissues have a relatively smaller change fold in expression level than those overlapping anchors of tissue-specific loops. Pairwise comparisons across four tissues are shown. Chromatin loops identified by hicDetectLoops were used for this analysis. **f**, 40-kb bins overlapping anchors of shared loops between tissues exhibit a significantly lower expression specificity index Tau value compared to those overlapping tissue-specific loops. Loops identified by hicDetectLoops were used for this analysis (Fig. 5b). All *P*-values reported were calculated with Wilcoxon signed rank exact tests.

We also performed the reciprocal analysis to see whether changes in gene expression corresponded to changes in subcompartments. In comparisons between pairs of tissues, all transcribed bins (24,038 testable 40-kb bins with CPM > 0.5) were assigned into three groups based on their changes of transcription level: (1) the down group, in which bins exhibited expression level decreases larger than 2 fold, (2) the up group, in which bins exhibited expression level increases larger than 2 fold, and (3) the stable group that included all other bins. By integrating subcompartment profiles, we observed that bins with increased subcompartment rank were slightly enriched in the up expression group, while bins with decreases in subcompartment rank were slightly enriched in the down group (see **Fig. 7b** for comparison between bud and leaf, and other five comparisons in Extended Data Fig. 11b). However, a large fraction of bins (e.g. 64.1-65.3% in the comparison between bud and leaf exhibited stable subcompartment ranks (rank change = 0) in all three groups. These results suggest that changes in gene expression are only associated with subcompartment patterning for a small subset of genomic regions, because most differentially transcribed bins remain unchanged subcompartments.

We extended these analyses to TADs, as opposed to compartments, and asked whether remodeling TAD structures related to differential gene expression between tissues. To do so, we performed a pairwise comparison of both the TAD profiles and the transcriptomes of the four pepper tissues. For simplicity, we divided TADs and TAD boundaries into two groups: conserved between tissues and tissue-specific. Based on TADs annotated by Arrowhead, we did not detect enrichment of differentially expressed genes for either TADs or boundaries in the tissue-specific group compared to the conserved group (Supplementary Table S16). However, we did find that conserved TAD boundaries are associated with a significantly lower change of expression level than tissue-specific boundaries, as measured by the absolute fold-change in expression level for each 40-kb bin (Wilcoxon rank-sum test *p* < 0.0025 for five comparisons; **Fig. 7c**). This pattern was not observed for TAD bodies (Extended Data Fig. 12a,b). Furthermore, when we studied the expression specificity index *Tau* value instead of fold-change in expression level by stratifying TADs and TAD boundaries by their stability across tissues, we observed that TAD boundaries shared between/across tissues were associated with a significantly smaller variation in expression level than those unique to a specific tissue (Wilcoxon rank-sum test *p* < 0.0014; **Fig. 7d**). As with fold-change, this is not observed for TAD bodies (Extended Data Fig. 12c,d). All of these observations were consistent in TAD annotations from TopDom (Extended Data Fig. 12e,f). Overall, these results suggest that TAD structures are associated with gene regulation in a way that is largely confined to genes in or near the TAD boundary regions.

Finally, we examined whether variation in chromatin loops is associated with changes in gene expression by comparing loops (based on hicDetectLoops inferred loops) that are shared in two or more tissues (5,728) and those unique to a single tissue (13,793). Similar to results for subcompartments and TAD boundaries, differentially expressed genes were not enriched for either loop group (Supplementary Table S17). However, we found that loops shared across tissues were associated with more stable expression level than tissue-specific loops, as shown by their fold changes in expression level (**Fig. 7e**) and their *Tau* values (**Fig. 7f**). These results paralleled those based on TAD boundaries. Taken together, our results suggest that 3D chromatin folding does not directly determine differentially gene regulation and expression, but rather creates an environment or framework consolidating other mechanisms controlling gene expression.

## Discussion

In this study, we have created a reference-grade genome assembly for *Capsicum annuum* (pepper) accession *CA59* and performed Hi-C to dissect patterns of 3D genome folding in four tissues. We have found that the pepper genome is predominantly partitioned into TADs and that a large proportion of TADs are likely formed by loop extrusion. We suggest that the diversity of TAD categories in plants can be reconciled by the underlying formation mechanisms whose prevalence may be related to genome size and transposable element content. We also provided evidence that the 3D genome influences the patterns of genomic structural variations in plants. And our integrated analysis of Hi-C profiles and transcriptomes across tissues demonstrated that 3D chromatin organization does not directly contribute to differential gene expression between tissues, but instead it does play roles in transcription stability.

On a global level, Hi-C contact maps derived from different tissues vary considerably. We have observed a distinct anti-diagonal pattern of contacts in leaf and bud Hi-C contact maps, but this pattern was much weaker or absent in pulp and placenta (Fig. 1a). This contrast may reflect different chromosome conformations. Indeed, the conformation contrasts may derive from tissues exhibiting differences in so-called Rabl or non-Rabl configuration of interphase nuclei. Such inter-tissue differences have been observed in other plant species ^68,69^. Additionally, we have observed an enrichment in the long-range (>20 Mb) and a depletion in the short-range (<20 Mb) interaction frequencies in the Hi-C contact maps of leaf and bud relative to those of pulp and placenta (Fig. 1c,d). This discrepancy may be a result of a difference in nuclear volume between different tissue cells ^70^. Surprisingly, although remarkable differences exist in global chromosome conformation between tissues, we still found subcompartments (Fig. 2f,h), TADs (Fig. 3c), and loops (Extended Data Fig. 9a) tend to be maintained across tissues. Further investigations will illuminate the role of global chromosomal morphology in local chromatin folding patterns ^71^.

Our data allowed inference of Hi-C subcompartments at multiple-levels. Using *Calder* and higher resolution (40 kb) Hi-C contact maps, we identified eight subcompartments that partition the A and B compartments. Interestingly, these inferred subcompartments are significantly correlated with transcription levels (Fig. 2e) and the distribution of genes versus TEs (Fig. 2f), suggesting that this classification is biologically meaningful. These subcompartments may also be associated with chromatin state, as shown previously in both animals and plants ^8,9,41^.

One striking finding in this study is that TADs in pepper are as conspicuous as those routinely annotated in animals. In other plant studies, TAD architecture is either less defined ^40,41^ or nearly absent ^39,72,73^. These observations suggest a relationship between genome size and TAD architecture in plants ^12,44^ - i.e. larger genomes tend to have more defined TADs. Given how important TEs are to plant genome size, we hypothesize that TEs play an important role in mediating the relationship between TAD architecture and genome size. Indeed, the most visible TADs in pepper Hi-C contact maps appear as long genomic segments with higher retrotransposon density than their flanking regions (Fig. 4f and Extend Data Fig. 8d), highlighting the potential role of retrotransposons in TAD formation. TE activity is a major driving force for genetic and phenotypic diversity in flowering plants ^74,75^. Recent studies have also shown that TEs contribute to divergence and rearrangement of 3D chromatin organization ^76,77^, suggesting they not only contribute to structural patterns of 3D genome organization but also play a mechanistic role in shaping chromatin structures. If TEs do participate in organizing the 3D structure constituting TADs, future work needs to investigate which TE features mediate this role, i.e., the relative roles of specific TE families, transcriptional activity, sequence motifs and epigenetic effects.

Loop extrusion and compartmentalization are two major mechanisms for TAD formation ^4^, which cooperate to establish the 3D organization of the genome ^55^. Similar to wheat ^42,43^, TAD boundaries in pepper are frequently demarcated by chromatin loops, whereas this pattern is much less frequent in tomato and maize, and rare in other plant species with smaller genome sizes ^41^, suggesting loop extrusion is more prevalent in large compared to smaller plant genomes. Even so, TADs in tomato and maize largely coincide with compartments as evidenced by the fact that most TAD boundaries (55% in tomato) overlap with compartment borders ^41^. This suggests that compartmentalization is more prevalent in other genomes, where TAD boundaries less frequently overlap with compartment boundaries. We demonstrate this difference by comparing our pepper annotations to tomato annotations, using the same tools. We found that only ∼11% of TADs in pepper coincide with subcompartments (i.e. exhibit ≥ 80% reciprocal overlap of segments), compared to ∼30% in tomato. Based on these observations, we hypothesized that loop extrusion tends to facilitate the formation of large repressed TADs that are well-defined in Hi-C maps, whereas compartmentalization tends to facilitate the formation of smaller, less well-defined active TADs. This might be a result of condensed chromatin in inactive TADs forcing segments to be in close proximity, which may not be the case in less compacted active TADs ^21^. The latter point is supported by the observation that active TADs possess a more homogeneous state of subcompartment in their bodies than inactive TADs (**Fig 4b** and Extended Data Fig. 8c). Altogether, results suggest both that the relative prevalence of two leading mechanisms of TAD formation varies considerably across plant species and that these dynamics may explain variation in TAD prevalence. At least one question remains unresolved, however, which is whether additional mechanisms are involved in TAD formation in plants. Previous studies have revealed that sequence features, like the physical structure of genes ^33^, functional noncoding sequences ^78^, and transposable elements or their activities ^14,77^, are associated with TAD structure or their formation. The idea that sequence content affects TAD formation is consistent with the fact that boundaries tend to be near genes (Fig. 4h) and many prominent TADs are enriched in retrotransposons (Fig. 4f) in pepper.

The cause-and-effect relationships between key features of 3D genome organization and gene transcription remains an issue of open debate. Our integrated analysis of major chromatin features (e.g. subcompartments, TADs, and chromatin loops) and transcriptomes across four tissues has revealed that changes in chromatin structures do not directly contribute to differential gene expression between tissues. This is evidenced by the observations that significantly differentially expressed genes are not enriched for genomic regions where changes in chromatin folding occurred between tissues. Additionally, differentially transcribed genomic regions always exhibit unchanged 3D genome folding. However, we have also observed that the stability of features of 3D genome organization are subtly associated with gene expression across tissues (Fig. 7). Combining previously published studies that investigated the relationship between 3D chromatin structures and gene expression on different cell types and evolutionary time scales ^16,18,20,33^, the literature is reaching a consensus that genome architecture does affect broad patterns of gene expression. Notably, restructuring TADs is associated with expression change for genes located at the boundaries but not for genes located inside them, which suggests TADs may act as an architectural framework for the establishment of long-range regulation of genes at their boundaries. Our results suggest that changes in chromatin folding are not a primary mechanism contributing to differential gene regulation and expression of single genes, which is consistent with recent studies in mammals as well as in *Drosophila*. Instead, these studies suggest that chromatin conformation provides a structural scaffold for the establishment of the regulatory environment ^22–25^.

The chromatin loop anchors we identified here are enriched for gene bodies. This is also observed in wheat ^42^ and many other large plant genomes ^41^, leading to the abundance of gene-to-gene loops in these genomes. This kind of loop could result from the interaction of gene islands dispersed across repetitive heterochromatic regions, potentially resulting in three dimensional clusters of genes that are otherwise dispersed in sequence space (Fig. 5e,f). Physically clustered genes are common in plants and tend to be co-regulated and possess similar functions ^60,79^. It is worth noting that the loops reported here are annotated based on Hi-C maps of bin size 10kb or larger, which may be somewhat different in their form and function from chromatin loops that are inferred at gene or kilobase scales (i.e. enhancer-promoter loops) using higher-resolution chromatin interaction maps ^26,28,39^.

Similar to observations in animals ^18,31,33^, breaks of chromosomal synteny in pepper, eggplant, potato, and tomato genomes preferentially occur at TAD boundaries (Fig. 6f). Such a pattern may be due to higher chromatin fragility at TAD boundaries and/or increased selective pressure against rearrangements that disrupt TAD integrity ^30,33,80^. In animals, TADs appear to behave as structural and functional units of the genome and can still be highly conserved between species separated by several million years ^5,33,81^. In contrast, TADs show almost no conservation when comparing distantly diverged plant species ^41^. This discrepancy could result from the different TAD properties between both systems, including structural (sequence determinants), mechanistic (formation mechanisms), and functional (i.e. constraints) features. Additionally, TAD boundaries are also characterized by depletion of structural variants and SNPs, similar to those observed in animals ^33,35,82–84^, consistent with the enrichment of evolutionary sequence conservation at such regions. Notably, these patterns are not observed in rice ^61^, suggesting that the functional implications of TADs might be as diverse between plant species as their putative mechanisms of formation. Integrating these different patterns and conflicting results into a coherent framework explaining their mechanistic origins and functional consequences will require high resolution annotation of chromatin structures between different systems and organisms.

In summary, our results suggest that TAD architecture in plants exhibits considerable diversity, especially when compared to animal systems. This diversity may be mediated by distinct formation mechanisms that vary according to genome size, gene content, TE compositions, or transcription state. Clarifying the mechanisms that lead to the observed diversity may help to understand the functional roles of TADs in different organisms and the relationship between genome function and 3D genome organization, such as whether high-order chromatin structures contribute to genome function or just a result of genome function. Additionally, understanding of principles of 3D genome folding, formation mechanisms, and sequence determinants of key chromatin features in plants would also facilitate sequence-based modeling and engineering of genome 3D architecture ^85,86^. Increasing Hi-C data from different plants and advancing genome capture technology will facilitate more and better comparative analyses. They will deepen our understanding of the principles of 3D genome folding, diversity across organisms, and the function of chromatin structures.

## Methods

### Plant materials and DNA sequencing

An elite pepper (*Capsicum annuum*) inbred line, designated as ‘CA59’, was used in this study due to its desirable agronomic characteristics, including high yield, broad-spectrum disease resistance, and abiotic stress tolerance ^46^. Seeds were germinated in soil in 72 cell plastic flats, placed in the greenhouse on February 2nd and August 6th. The seedlings were grown in a greenhouse under normal conditions in Guangzhou, China (23.1291° N, 113.2644° E).

Thirty-day-old fresh leaves harvested from a single individual plant were used for DNA extraction and sequencing. For BGI (Beijing Genomics Institute) short-read sequencing, DNA was extracted from about 2 g leaves using a modified cetyltrimethylammonium bromide (CTAB) method ^87^. A sequencing library with an insert size of 350 bp was prepared using the VAHTS Universal DNA Library Prep Kit (Vazyme, Nanjing, China). Quality assessment of the library assessing DNA quantity, purity, and size range was conducted using Agilent Bioanalyzer 2100 (Agilent Technologies, Santa Clara, CA). The library was sequenced on the MGI-SEQ 2000 sequencing platform to produce pair-end sequence data (2 × 150 bp). For Pacific Biosciences (PacBio) sequencing, extraction of high-molecular-weight DNA was carried out following the protocol previously described. About 10 μg of genomic DNA was used to prepare template libraries of 30-40 kb using the BluePippin Size Selection system (Sage Science, USA) following the manufacturer’s protocol (Pacific Biosciences, USA). The libraries were sequenced on the PacBio SEQUEL II platform with three SMRT flow cells.

### Transcriptome sequencing

We generated long-read full-length transcriptomes for five tissues, including leaves, flower buds, placentas, roots, and pulp, using the PacBio isoform sequencing (Iso-seq) platform. Using the SMRTlink 8.0 pipeline (https://www.pacb.com/support/software-downloads/), we assembled between 35,257 and 50,237 transcripts across these five tissues (Supplemental Table S5). These transcripts were used for gene annotation.

Accordingly, we generated RNA-seq data for the same tissues, each with three biological replicates (Supplementary Table S7). Samples used for RNA extraction were pooled from 5 individual plants and total RNA was extracted using Trizol reagent following the manufacturer’s recommendations (Invitrogen, CA, USA). RNA purity and integrity were assessed using NanoDrop 2000 spectrophotometer (NanoDrop Technologies, Wilmington, DE, USA) and Bioanalyzer 2100 system (Agilent Technologies, CA, USA). RNA contamination was assessed using 1.5% agarose gel electrophoresis. A total of 1 μg of RNA per sample was used as the input material for library preparation. The mRNA was purified from the total RNA using poly-T oligo-attached magnetic beads. Sequencing libraries were generated from the purified mRNA using the V AHTS Universal V6 RNA-seq Library Kit for MGI (V azyme, Nanjing, China) following the manufacturer’s recommendations with unique index codes. The size of the resulting library was assessed using Qubit 3.0 Fluorometer (Life Technologies, Carlsbad, CA, USA) and Bioanalyzer 2100 system (Agilent Technologies, CA, USA). Subsequently, sequencing was performed on the MGI-SEQ 2000 platform by Frasergen Bioinformatics Co., Ltd. (Wuhan, China).

### In situ Hi-C library construction protocol, sequencing, and data processing

We generated in situ Hi-C data for four tissues, including leaves, placentas, pulp, and buds. Hi-C libraries of two biological replicates (Supplementary Table S18) were constructed for each group according to the protocol established by Rao et. al. ^8^. Briefly, about 2 g of plant material was cut into 1- to 2-mm strips, which were fixed with 2% final concentration fresh formaldehyde in NIB buffer (20 mM HEPES, pH 8.0, 250 mM sucrose, 1 mM MgCl2,5mM KCl, 40% (v/v) glycerol, 0.25% (v/v) Triton X-100, 0.1 mM PMSF, and 0.1% (v/v) β-mercaptoethanol) at 4°C for 45 min in a vacuum. Formaldehyde was added at a final concentration of 0.375 M glycine under vacuum infiltration for an additional 5 min. The samples were washed twice in ice-cold water. The clean samples were frozen in liquid nitrogen and then ground to a powder and resuspended in NIB buffer. The solution was then filtered through one layer of Miracloth. The nuclei isolated from these tissues were lysed with 0.1% (w/v) final concentration SDS at 65°C for 10 min and then SDS molecules were added using Triton X-100 at a 1% (v/v) final concentration. The DNA in the nuclei was then digested by adding 200U MboI (NEB) and incubating the samples at 37°C for 2 hr. Restriction fragment ends were labeled with biotinylated cytosine nucleotides by biotin-14-dCTP (TriLINK). Blunt-end ligation was carried out at 16°C overnight in the presence of 50 Weiss units of T4 DNA ligase. After ligation, the cross-linking was reversed by 200 μg/mL proteinase K (Thermo) at 65°C overnight. DNA purification was achieved through QIAamp DNA Mini Kit (Qiagen) according to the manufacturer’s instructions. Purified DNA was sheared to a length of ∼400 bp. Point ligation junctions were pulled down by Dynabeads® MyOne™ Streptavidin C1(Thermofisher) according to manufacturer’s instructions. The Hi-C library for Illumina sequencing was prepped using the NEBNext® Ultra™ II DNA library Prep Kit for Illumina (NEB) according to manufacturer’s instructions. Fragments between 400 and 600 bp were paired-end sequenced on the Illumina HiSeq X Ten platform (San Diego, CA, United States) at 150 PE mode.

Raw Hi-C reads were cleaned using Trimmomatic (version 0.38) ^88^. Hi-C contact maps were constructed using both Juicer ^50^ and HiCExplorer version 3.53 ^11^ pipeline. A detailed description of the processing is provided in Supplementary Methods. Quality and reproducibility of the Hi-C data were assessed using QuASAR-Rep analysis (3DChromatin-ReplicateQC version 0.0.1) ^51^ and Pearson correlation analysis with hicCorrelate command in HiCExplorer.

### Genome assembly and assessment

The genome size of *Capsicum annuum* was estimated using k-mer frequency distribution generated from 353.90 Gb cleaned BGI short reads. GCE (Genome Characteristics Estimation) software (ftp://ftp.genomics.org.cn/pub/gce) was used to calculate 17-mer frequency distribution and estimate the genome size with modified parameters: -m 1 -D 8 -b 0 -H 1. This estimated genome size is about 2.950 Gb and the heterozygous rate is 0.23%.

We obtained about 451 Gb clean PacBio long reads (∼150× genomic coverage) from three SMRT flow cells with a subread N50 of 28,351 bp. Before assembly, we filtered out short-length PacBio raw reads and only retained the top 200 Gb longest reads (subread N50 is 39,818 bp, ∼66 × genomic coverage) for further correction using MECAT2 version 2.1 ^89^. The corrected PacBio reads were then trimmed and assembled with CANU version 2.0 ^90^. The initial contig assembly was polished through three iterations using Pilon version 1.23 ^91^ with 120-fold BGI short reads. Finally, we used the Juicer, Juicerbox, 3D-DNA pipeline ^50,92,93^ with a combination of Hi-C data from two tissues, flower bud and leaf, totaling 415.16 Gb, corresponding to ∼135X genome coverage, for genome scaffolding and manual correction. A detailed summary of above assembly processes and the parameters are provided in Supplementary Methods.

For assembly assessment, the gene-space completeness was estimated using BUSCO (v3.0.2) with the Embryophyta odb9 dataset (*n* = 1440) ^94^. Synteny analysis was performed using Minimap2 ^95^ and pafr (https://github.com/dwinter/pafr) was used to make a dotplot between scaffolds and the existing *Capsicum annuum* cv Zunla-1 genome ^96^ and three more distantly solanaceous species, including tomato, potato, and eggplant.

### Transposable elements (TE) and gene annotation

Transposable elements were annotated with the Extensive *de-novo* TE Annotator (EDTA) pipeline (version 1.9.6) ^97^. Gene models were constructed using the MAKER pipeline ^98^, performed in three iterations. The EDTA repeat library, Iso-seq full-length transcripts from five tissues, and gene models from the *Zunla*-1 assembly ^47^ were used as evidence-based resources for MAKER to build gene models. Additionally, RNA-seq reads from five tissues were mapped to the *CA59* genome using HISAT2 ^99^. An improved transcriptome was then constructed using StringTie ^100^. The predicted new transcripts were merged with the MAKER gene models to produce the final gene/transcript set. A detailed description of the annotation is provided in Supplementary Methods.

### Compartment and subcompartment calling

To identify the A and B compartments, we first adapted the method as described previously ^1,19^. First, the observed/expected matrices were calculated with normalized and corrected (ICE) interaction matrices for each chromosome at 500-kb resolution. Next, Pearson correlation and covariance matrices were computed on the observed/expected matrices. Third, PCA eigenvectors were calculated with the covariance matrices and the first principal component (PC1) was used to assign the A and B compartments according to the direction of the eigenvalues which were manually adjusted by the gene and TE density. All these steps were processed using the HiCExplorer tool ^11^. This method is capable of identifying the A and B compartments globally when working on Hi-C maps at the 500-kb resolution, whereas it failed to identify the A and B compartments consistently when using Hi-C maps at a relatively higher resolution (e.g., 40-kb).

To further determine compartments at a finer resolution, we utilized Calder ^13^ which is capable of inferring multiple sub-compartments using low-resolution Hi-C data. To do this, the HiCExplorer interaction matrices at 40-kb resolution were first transformed to a square format and were then imported into the R package, *BNBC* ^56^, for normalization and batch correction across tissues and replicates. The resulting matrices were converted into a three-column format which is required as input for Calder. We ran Calder with the default parameters which call eight sub-compartments, of which, 4 sub-compartments (designated as A.1.1, A1.2, A2.1, and A2.2) belonging to the A compartment and the other 4 (designated as B1.1, B1.2, B2.1, and B2.2) belonging to the B compartment. To identify genomic regions that exhibit A/B compartments or sub-compartment switching, we performed analyses by considering: 1) changes in one replicate, and 2) changes supported in both replicates.

### TAD calling and classification

TADs were annotated using three distinct tools, including HiCFindTADs ^11^, TopDom ^101^, and Arrowhead ^8^. The latter two were chosen because of their top performance in a previous benchmarking study ^58^. To evaluate the consistency of TAD calls among tools, we first applied these three tools to leaf Hi-C interaction matrices at 40 kb resolution. We measured the overlap of TADs, TAD boundaries, and genome coverage among TAD sets derived from different callers. A reciprocal overlap threshold of >80% was used to define TADs that are consistently identified by the different tools used. TAD boundaries were considered as overlapping if they were less than one bin size, i.e., 40 kb, apart.

To assess the reproducibility of TAD calls across samples (tissues and replicates), we chose to use TopDom for TAD calling, which is compatible with both the continuous TAD distribution manner in the pepper genome and also with the output of the BNBC program ^56^ that we used to normalize and correct Hi-C matrices. TAD calls from another two tools were also compared as needed. Hierarchical clustering analysis was performed based on the Jaccard distance of the genome coverage of overlapping TADs across samples.

The hierarchical clustering of TADs was performed using the *hclust* function with the ‘average’ method in R version 4.0.4. The similarity was calculated by taking sub-compartment signalings (i.e., B2.2:0.125; B2.1:0.25; B1.2:0.375, B1.1:0.5; A2.2:0.625; A2.1:0.75; A1.2:0.875; A1.1:1) for each bin that spans TAD regions as the input. A heatmap was constructed using the *heatmap*.*2* function in R.

Hi-C contact maps were displayed using hicPlotMatrix from the HiCExplorer tool and the online JuiceBox tool https://aidenlab.org/juicebox/.

### Synteny breaks

Identification of evolutionary synteny breaks between the query genome and the reference genome was conducted as previously described ^33^ with minor modifications to fit the comparison of large plant genomes. For example, whole-genome alignments were conducted using minimap2 version 2.17 in this study. Synteny breaks were identified from pairwise comparisons among four distantly related Solanaceae genomes, including pepper, tomato, eggplant, and potato when using each of them except eggplant as the reference.

To quantify the distribution of evolutionary synteny breaks along the TADs, we adapted the method described in Krefting et al. 2018. ^18^. We noted that TAD boundaries are enriched for evolutionary sequence conservation which might result in an enrichment of synteny breaks identified in such regions. Therefore, we further normalized the observed distribution using the rate of the conserved alignable sequences along the TAD bodies. A significance test was performed by generating 100 sets of random synteny breaks as a background control in each comparison.

### Genomic variants calling and patterns around TAD boundaries

We used a custom assembly-based pipeline to identify SNPs and deletions from five closely related genomes, including two within-species accessions, *CM334* and *Zunla-1*, a wild progenitor, *glabriusculum*, as well as two closely related species, *C. chinense* and *C. baccatum*, relative to the *CA59* genome. We excluded insertions for downstream analysis due to assembly quality issues.

The assembly-based structural variant detection pipeline includes four key steps: (1) The query genomes are aligned against the reference genome using minimap2 version 2.17 ^95^. (2) The resulting Axt alignment files from step 1 are used to build long genome alignment chains (i.e., connect alignments if they are close enough) using axtChain. The chain files were sorted and merged into a single file using chainMergeSort as necessary. Next, a filtering procedure was applied to remove the low-quality chains using chainPreNet and the remaining chains were then used to determine alignment nets by running chainNet. Finally, synteny information was added using netSyntenic. (3) SV calling for each pairwise comparison was performed using our custom Perl scripts based on the final syntenic format file from step 2. The SV output includes insertion, deletion, tandem duplication, inversion, and complex SVs. (4) Population-scale genotyping of SVs was performed using custom Perl scripts.

To measure the relative abundance of genomic variants around TAD boundaries, we used a sliding window approach with a bin size of 40 kb and a step size of 5-kb to generate an observed/expected matrix for 500 kb on each side. We defined the expected abundance under the null hypothesis that genomic variants are homogeneously distributed along the genome. The custom Perl scripts for this pipeline are available at https://github.com/yiliao1022/.

### Loop calling

Chromatin loops were annotated using the hicDetectLoops tool from HiCExplorer version 3.5.3 ^11^. To obtain denser Hi-C interaction matrices, we combined Hi-C data from two replicates for each tissue. Hi-C interaction matrices were normalized using the Knight-Ruiz (KR) method. Our visual inspection of Hi-C contact maps shows extensive loops that can span over several megabases. Therefore, we called loops from Hi-C interaction matrices at multiple resolutions, including 10kb, 15kb, 20kb, and 25kb. Loops identified from all resolutions were merged within 25 kb to produce the final loop set using hicMergeLoops from HiCExplorer version 3.5.3. Additionally, we used Mustache ^59^ to call loops with the juicer Hi-C interaction matrices for comparison. Parameters used for loop calling were provided in Supplementary Table S13. We used the *Intersect* function from pgltools version 2.2.0 ^102^ with the parameter: ‘-d 25kb’ to determine if loops are shared between tissues.

### Overlap between TADs with compartments and loops

To determine the extent of overlapping between TADs and *Calder*-inferred sub-compartments, we adapted a reciprocal overlap threshold of >80%. Since TADs and sub-compartments annotated with Hi-C matrices of different resolutions vary considerably in sizes, we performed multiple pairwise comparisons between TADs and sub-compartments inferred with different conditions. TADs were annotated using HiCExplorer, TopDom, and Arrowhead for 40-kb, and 100-kb resolution matrices, with an additional call for 10-kb resolution matrices using HiCExplorer. Subcompartments were annotated at 40-kb and 100-kb resolution matrices.

To determine the frequency of TADs that are demarcated by loops, we constructed a *PGL* file by pairing TAD boundaries sequentially. Then, we used the *intersect* function in the pgltools version 2.2.0 tool to identify the overlapping of TADs and loops--that is, two boundaries of a TAD coincide with the two anchors of a loop.

### Expression patterns and differential expression analysis

RNA-seq data from five tissues were used to broadly characterize the expression patterns for each gene and 40-kb bin. We measured two properties of the expression pattern: expression level fold change between tissues and tissue specificity. Tissue specificity was calculated using the formula tau=sum (1-r_i_)/(n-1) ^106^, where r_i_ represents the ratio between the expression level in sample i and the maximum expression level across all tissues, and n represents the total number of tissues. The expression level for each gene/transcript was quantified in normalized TPM (Transcript Per Million) and FPKM (Reads Per Kilobase of transcript per Million reads mapped) values using FeatureCounts ^103^ and a custom R script. We obtained the fold change of expression level and differentially expressed genes/40-kb bins between tissues using Limma-Voom ^67,105^. For 40-kb bins, the tau value was computed based on the CPM (counts per million) values derived from the Limma-Voom analysis. This value ranges from 0 to 1, with higher values indicating greater variations in expression level across tissues and thus higher tissue specificity. All expression values were averaged among three replicates.

Alternatively, to facilitate integrated analysis of expression data and higher-order chromatin structures that are annotated based on genomic bins of a fixed size (e.g., 40 kb and 100kb), we calculated the number of reads per bin (coverage tracks) from RNA-seq alignments to represent the expression level for corresponding bins. Coverage tracks were generated using the bamCoverage tool from deepTools version 3.3.0 ^104^ with the following parameters: ‘--binSize 40000 --minMappingQuality 30 --outFileFormat bedgraph’. The expression level for each bin was normalized in CPM (counts per million).

### Integrated analysis of Hi-C profiles and transcriptomes

To assess the functional relationship between 3D genome organization and gene expression, we first ask whether changes in subcompartments correspond to changes in expression. To do this, the genome, measured in bins (e.g., 40-kb), was partitioned into three groups based on subcompartment switching between tissues: 1) the ‘down’ bins in which subcompartments transitioned for at least 1 rank from A1.1 to B2.2, 2) the ‘up’ bins in which subcompartments transitioned for at least 1 rank in reverse order, and 3) the ‘stable’ bins in which subcompartment unchanged. For this classification, we generated two sets of data: 1) switching only considered in one replicate and 2) switching was supported in both replicates. Then, we compared the number of differentially expressed genes/bins among groups and the log fold change of expression level for genes/40-kb bins that overlap with each group.

Second, we tested whether changes in TAD structures correspond to changes in expression. To do this, TADs and TAD boundaries were divided into two categories in each pairwise comparison between tissues: 1) tissue-specific and 2) conserved between tissues. We then compared the number of differentially expressed genes/bins overlapped with each group and the absolute fold change of expression level for genes/40-kb bins that overlapped with each group. Next, we stratified TADs and TAD boundaries by their stability across tissues. For example, using Arrowhead TAD annotation, we identified 3,105 TAD boundaries that are specific to only tissue, and 1,373, 881, and 734 are shared in 2, 3, and 4 tissues, respectively. We correlated this classification to the tau values. TAD annotated by Arrowhead and TopDom using Hi-C maps at 40-kb resolution were used for the above analyses.

We also tested whether changes in loops correspond to changes in transcription. As with TADs, loops were divided into two groups in each comparison between tissues: 1) shared between tissues, and 2) tissue-specific. We then compared the number of differentially transcribed bins that overlap with the anchors of loops in each group, and the absolute fold change in expression level for 40-kb bins that overlapped with the loop anchors in each group. Also, for simplicity, loops merged from all four tissues were divided into two categories: 1) tissue-specific, and 2) conserved across tissues (i.e. shared in two or more tissues). We correlated this classification to the tau values calculated for 40-kb bins. Loops identified using hicDetectLoops were used for this analysis.

## Data availability

The data that support the findings of this study have been deposited into CNGB Sequence Archive (CNSA) of China National GeneBank DataBase (CNGBdb) with accession number CNP0001129.

## Code availability

All scripts that reproduce analyses from the manuscript are available at GitHub (https://github.com/yiliao1022/Pepper3Dgenome).

## Acknowledgments

This work was supported by the National Natural Science Foundation of China (32070331, 32072580), the National Key Research and Development Program (2018YFD1000800), the Guangdong Basic and Applied Basic Research Foundation (2020A1515011396). This work was jointly funded by National Institutes of Health (R01GM123303), National Science Foundation (IOS-1656260), and start-up funding from the University of California, Irvine to J.J.E. We are grateful to Shujun Ou and Hongru Wang for critical reading of the manuscript.

## Author contributions

Yi L., J.J. E., and C.C. jointly supervised this work; Yi L. conceived of the presented ideas; C.C. carried out the experiments and collected the data; J.W. assembled and annotated the genome; Yi L. analysed the data and wrote the manuscript with input from J.W. and C.C.; J.J.E., and B.G. reviewed and edited thoroughly the manuscript; all authors discussed the results and commented on the manuscript.

## Competing interests

The authors declare no competing interests.

## Notes

### Competing Interest Statement

The authors have declared no competing interest.

