## Supplemental Material for "The 3D architecture of the pepper (*Capsicum annum*) genome and its relationship to function and evolution"

### Extended Data Figures

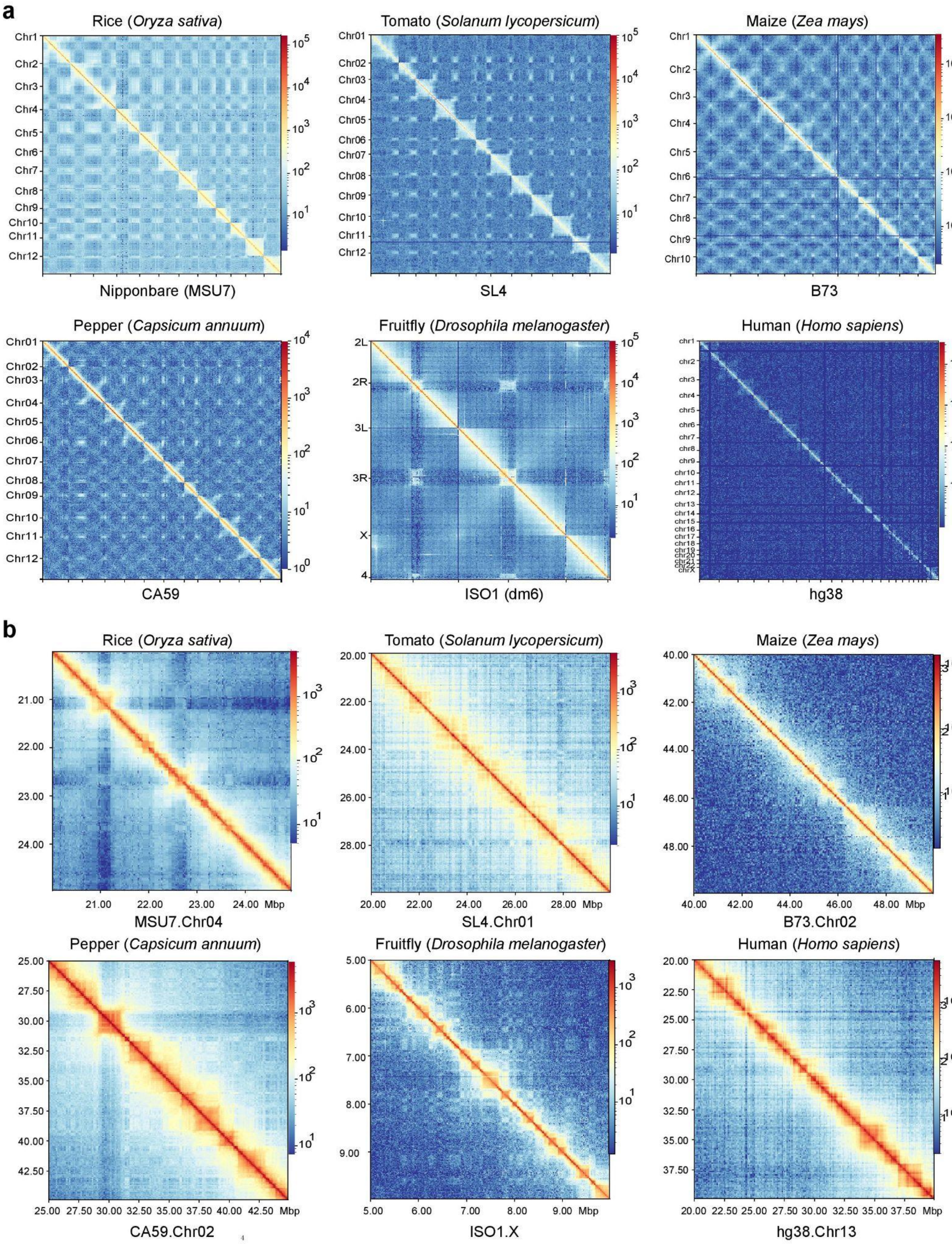

**Extended Data Fig. 1 | A preliminary visual inspection of Hi-C heatmaps in rice, tomato, maize, pepper, fruit fly, and human**. **a,** Genome-wide Hi-C heatmaps. Hi-C map resolution for each species: rice, 100 kb; tomato, 100 kb; maize, 500 kb; pepper, 500 kb; *Drosophila*, 100 kb; human 100 kb. **b,** TADs or similar structures (i.e. appear as clearly visible squares in the Hi-C maps) shown on example regions for each species using higher resolution Hi-C maps. Resolution: rice, 10 kb; tomato, 40 kb; maize, 100 kb; pepper, 100 kb; *Drosophila*, 5 kb; human 40 kb. Published Hi-C data used to generate the Hi-C maps can be found in Supplementary Table S19.

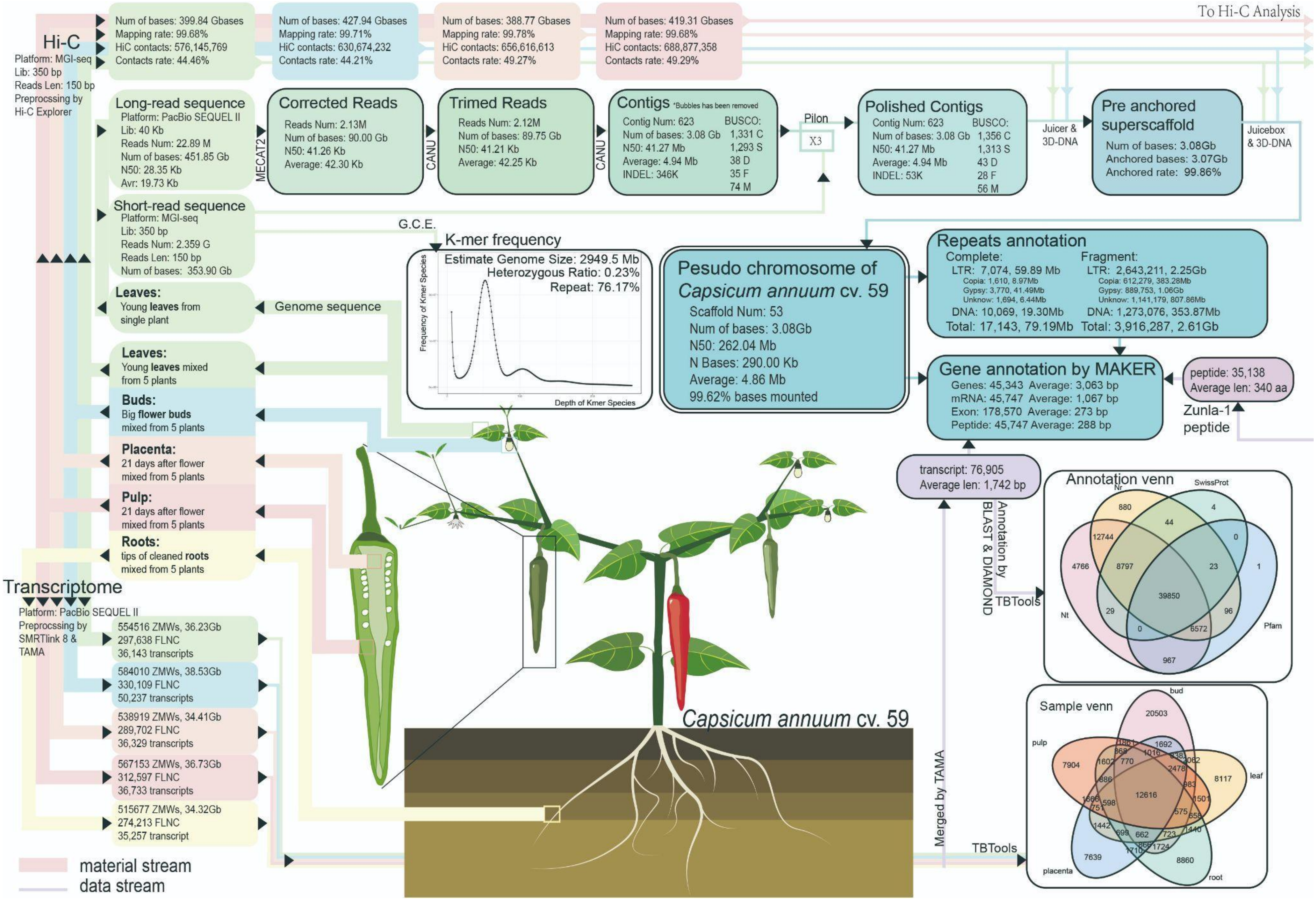

Hi-C
Platform: MGI-seq
Lib: 350 bp
Reads Len: 150 bp
Preprocssing by Hi-C Explorer
Num of bases: 399.84 Gbases
Mapping rate: 99.68%
HiC contacts: 576,145,769
Contacts rate: 44.46%
Num of bases: 427.94 Gbases
Mapping rate: 99.71%
HiC contacts: 630,674,232
Contacts rate: 44.21%
Num of bases: 388.77 Gbases
Mapping rate: 99.78%
HiC contacts: 656,616,613
Contacts rate: 49.27%
Num of bases: 419.31 Gbases
Mapping rate: 99.68%
HiC contacts: 688,877,358
Contacts rate: 49.29%
To Hi-C Analysis
Long-read sequence
Platform: PacBio SEQUEL II
Lib: 40 Kb
Reads Num: 22.89 M
Num of bases: 451.85 Gb
N50: 28.35 Kb
Avr: 19.73 Kb
MECAT2
Corrected Reads
Reads Num: 2.13M
Num of bases: 90.00 Gb
N50: 41.26 Kb
Average: 42.30 Kb
CANU
Trimed Reads
Reads Num: 2.12M
Num of bases: 89.75 Gb
N50: 41.21 Kb
Average: 42.25 Kb
CANU
Contigs *Bubbles has been removed
Contig Num: 623
Num of bases: 3.08 Gb
N50: 41.27 Mb
Average: 4.94 Mb
INDEL: 346K
BUSCO:
1,331 C
1,293 S
38 D
35 F
74 M
Pilon
X3
Polished Contigs
Contig Num: 623
Num of bases: 3.08 Gb
N50: 41.27 Mb
Average: 4.94 Mb
INDEL: 53K
BUSCO:
1,356 C
1,313 S
43 D
28 F
56 M
Juicer & 3D-DNA
Pre anchored superscaffold
Num of bases: 3.08Gb
Anchored bases: 3.07Gb
Anchored rate: 99.86%
Juicebox & 3D-DNA
Short-read sequence
Platform: MGI-seq
Lib: 350 bp
Reads Num: 2.359 G
Reads Len: 150 bp
Num of bases: 353.90 Gb
G.C.E.
K-mer frequency
Estimate Genome Size: 2949.5 Mb
Heterozygous Ratio: 0.23%
Repeat: 76.17%
Frequency of Kmer Species
Depth of Kmer Species
Pesudo chromosome of Capsicum annuum cv. 59
Scaffold Num: 53
Num of bases: 3.08Gb
N50: 262.04 Mb
N Bases: 290.00 Kb
Average: 4.86 Mb
99.62% bases mounted
Repeats annotation
Complete:
LTR: 7,074, 59.89 Mb
Copia: 1,610, 8.97Mb
Gypsy: 3,770, 41.49Mb
Unknow: 1,694, 6.44Mb
DNA: 10,069, 19.30Mb
Total: 17,143, 79.19Mb
Fragment:
LTR: 2,643,211, 2.25Gb
Copia: 612,279, 383.28Mb
Gypsy: 889,753, 1.06Gb
Unknow: 1,141,179, 807.86Mb
DNA: 1,273,076, 353.87Mb
Total: 3,916,287, 2.61Gb
Gene annotation by MAKER
Genes: 45,343 Average: 3,063 bp
mRNA: 45,747 Average: 1,067 bp
Exon: 178,570 Average: 273 bp
Peptide: 45,747 Average: 288 bp
peptide: 35,138
Average len: 340 aa
Zunla-1 peptide
Leaves:
Young leaves from single plant
Genome sequence
Leaves:
Young leaves mixed from 5 plants
Buds:
Big flower buds mixed from 5 plants
Placenta:
21 days after flower mixed from 5 plants
Pulp:
21 days after flower mixed from 5 plants
Roots:
tips of cleaned roots mixed from 5 plants
transcript: 76,905
Average len: 1,742 bp
Annotation by BLAST & DIAMOND
TBTools
Annotation venn
Nr
SwissProt
Nt
Pfam
880
4
44
12744
0
4766
8797
23
1
39850
29
96
0
6572
967
Transcriptome
Platform: PacBio SEQUEL II
Preprocssing by SMRTlink 8 & TAMA
554516 ZMWs, 36.23Gb
297,638 FLNC
36,143 transcripts
584010 ZMWs, 38.53Gb
330,109 FLNC
50,237 transcripts
538919 ZMWs, 34.41Gb
289,702 FLNC
36,329 transcripts
567153 ZMWs, 36.73Gb
312,597 FLNC
36,733 transcripts
515677 ZMWs, 34.32Gb
274,213 FLNC
35,257 transcript
Capsicum annuum cv. 59
Merged by TAMA
TBTools
Sample venn
bud
pulp
leaf
placenta
root
20503
7904
8117
7639
8860
12616
material stream
data stream

**Extended Data Fig. 2 | *De novo* sequencing, assembling, and annotation of the *CA59* genome based on PacBio long reads and chromosome conformation capture (Hi-C). (1)** For genome sequencing, we collected 451.85 Gb PacBio long reads, 353.9 Gb short reads, and 415 Gb Hi-C data (combined from a leaf and a bud sample). DNA was extracted from a single individual for DNA sequencing except for Hi-C experiments (see below). (**2**) For assembling, we started by selecting 200 Gb, the longest PacBio reads. This subset of reads were corrected using MECAT2, and further trimmed and assembled using CANU version 2.0. The draft assembly was then polished by short reads three rounds using Pilon version 1.23. Finally, chromosome conformation capture (Hi-C) was used to scaffold the contigs using the Juicer, JuiceBox, 3D-DNA pipeline. More details can be found in Methods and Supplementary methods. (**3**) For gene annotation, we collected PacBio Iso-seq sequencing data from 5 tissues, including leaf, bud, pulp, placenta, and root. Each tissue sample was harvested and merged from 5 individual plants. Gene models were predicted using the MAKER pipeline, integrating evidence including 5 tissues full-length transcript isoforms (built by SMRTlink8.0) obtained from the PacBio Iso-seq method, and gene models from a previous pepper accession, Zunla-1. Transposable elements were predicted by the EDTA pipeline. (4) For the architecture of 3D genome inference, we collected Hi-C data from 4 tissues, including leaf, bud, pulp, and placenta, each with two biology replicates.

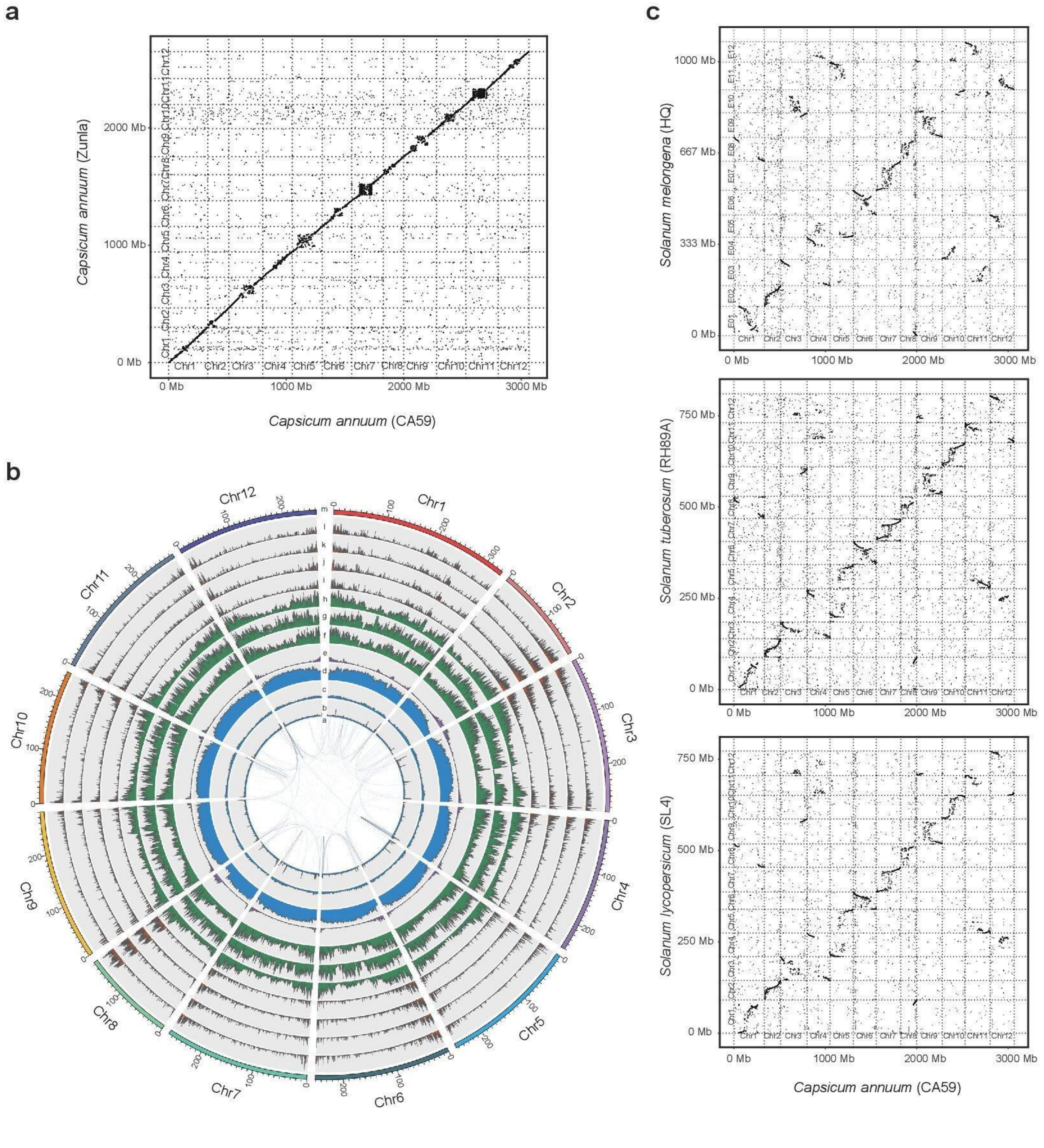
a
Capsicum annuum (Zunla)
Capsicum annuum (CA59)
b
c
Solanum melongena (HQ)
Solanum tuberosum (RH89A)
Solanum lycopersicum (SL4)
Capsicum annuum (CA59)

**Extended Data Fig.3 | Genomic features of the *CA59* genome and its synteny with other closely related genomes**. **a,** Syntenic dotplot between the *Capsicum annuum* cv. CA59 and cv. Zunla-1 assemblies. **b,** A circos diagram showing distribution of genomic features. a-e: intra-genome duplications, simple repeats, DNA transposons, LTR retrotransposons, Genes; f-h: SNPs, InDels, SVs (>50bp) identified from five closely related genomes (see **Fig. 6a**); i-l: transcription profiles in leaf, bud, pulp, and placenta. **c,** Syntenic dotplot between the *Capsicum annuum* cv. CA59 assembly and genomes of three more distantly related Solanaceae species, including eggplant (*S. melongena*), potato (*S. tuberosum*), and tomato (*S. lycopersicum*).

a

Intact LTR elements identified per haploid genome in some Solanaceae plants and maize

| Species | Genome size (Mb) | Num. of intact elements | Genome occupancy |
|---|---|---|---|
| Pepper | 3,077.7 | 7,074 | 1.95% (59.89 / 3077.7 Mb) |
| Tomato | 782.5 | 3,413 | 3.31% (25.9 / 782.5 Mb) |
| Eggplant | 1,073.1 | 4,208 | 3.40% (36.5 / 1,073.1 Mb) |
| Potato | 810.1 | 3,357 | 3.28% (26.6 / 810.1 Mb) |
| Maize | 2,131.8 | 51,980 | 23.92% (509.9 / 2,131.8 Mb) |

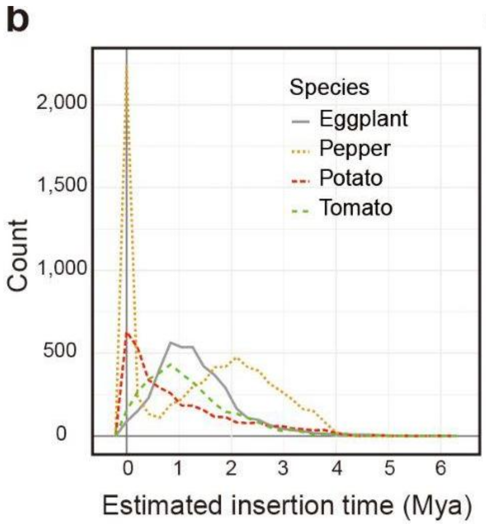


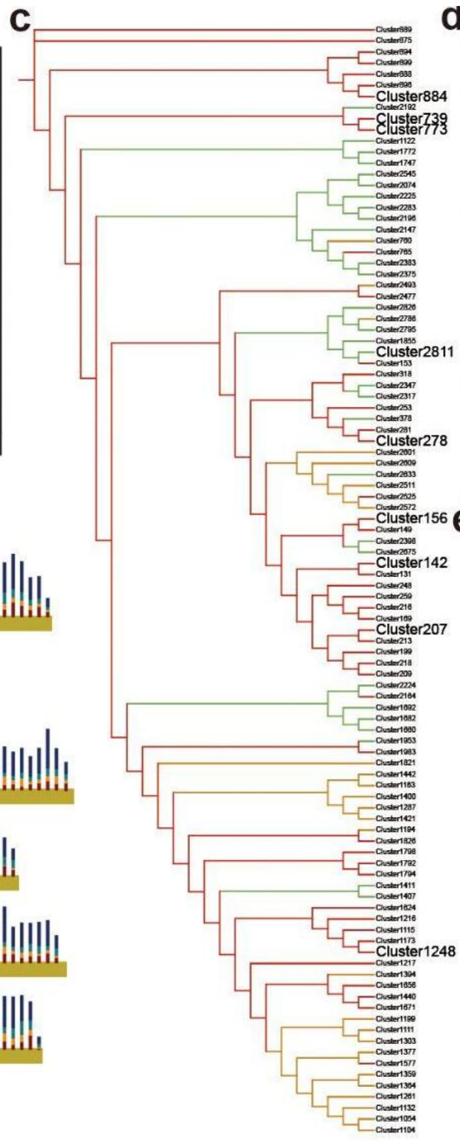


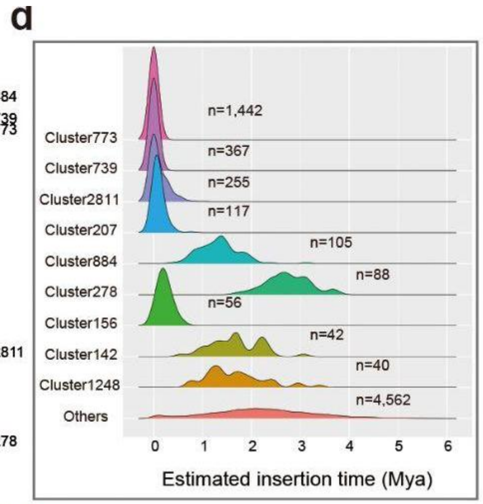


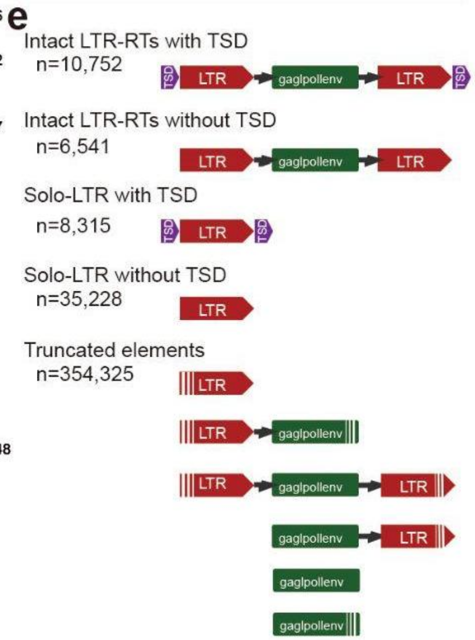


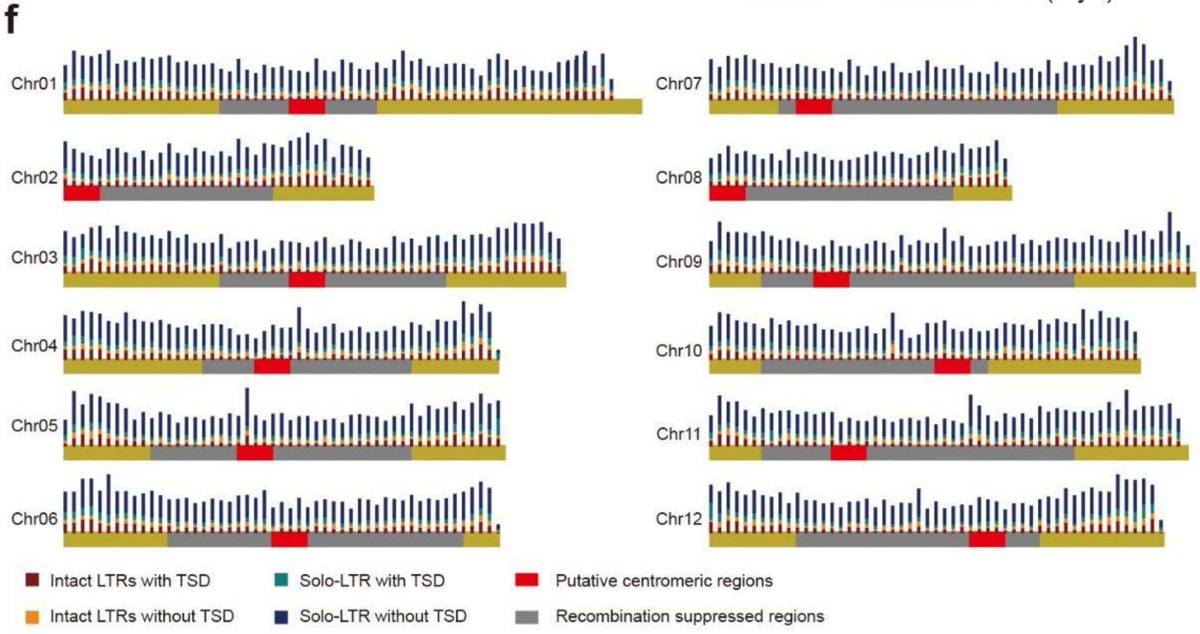

**Extended Data Fig. 4 | Analysis of LTR-RTs in the CA59 genome assembly.** **a,** Sequence occupancy of the intact LTR retrotransposons (LTR-RTs) in the genome of the four Solanaceae species, including pepper (CA59), tomato (SL4), eggplant (HQ), and potato (RH89A), together with maize (B73). **b,** Distribution of the estimated insertion times of intact LTR-RTs in the genome of each Solanaceae species. **c,** Phylogenetic relationship of the top 50 largest LTR-RT families in the *CA59* genome. Red branches indicate the *gypsy* family, green indicate the *copia* family, and orange undetermined family. **d,** Estimated insertion time of the top 9 largest LTR-RT families in the *CA59* genome. Of them, five families, including cluster 773, 739, 2811, 207, and 156, totaling 2,102 copies, with estimated insertion times almost near to zero, indicating they derived from very recent bursts of retroposition. **e,** Schematic representation of the structure of LTR-retrotransposon elements. **f,** Distribution of intact LTR-RTs, solo-LTRs, and fragmented segments along the chromosomes. Categories were summarized for each 5-Mb window. The centromere positions and recombination suppressed regions are taken from a previous work [47].

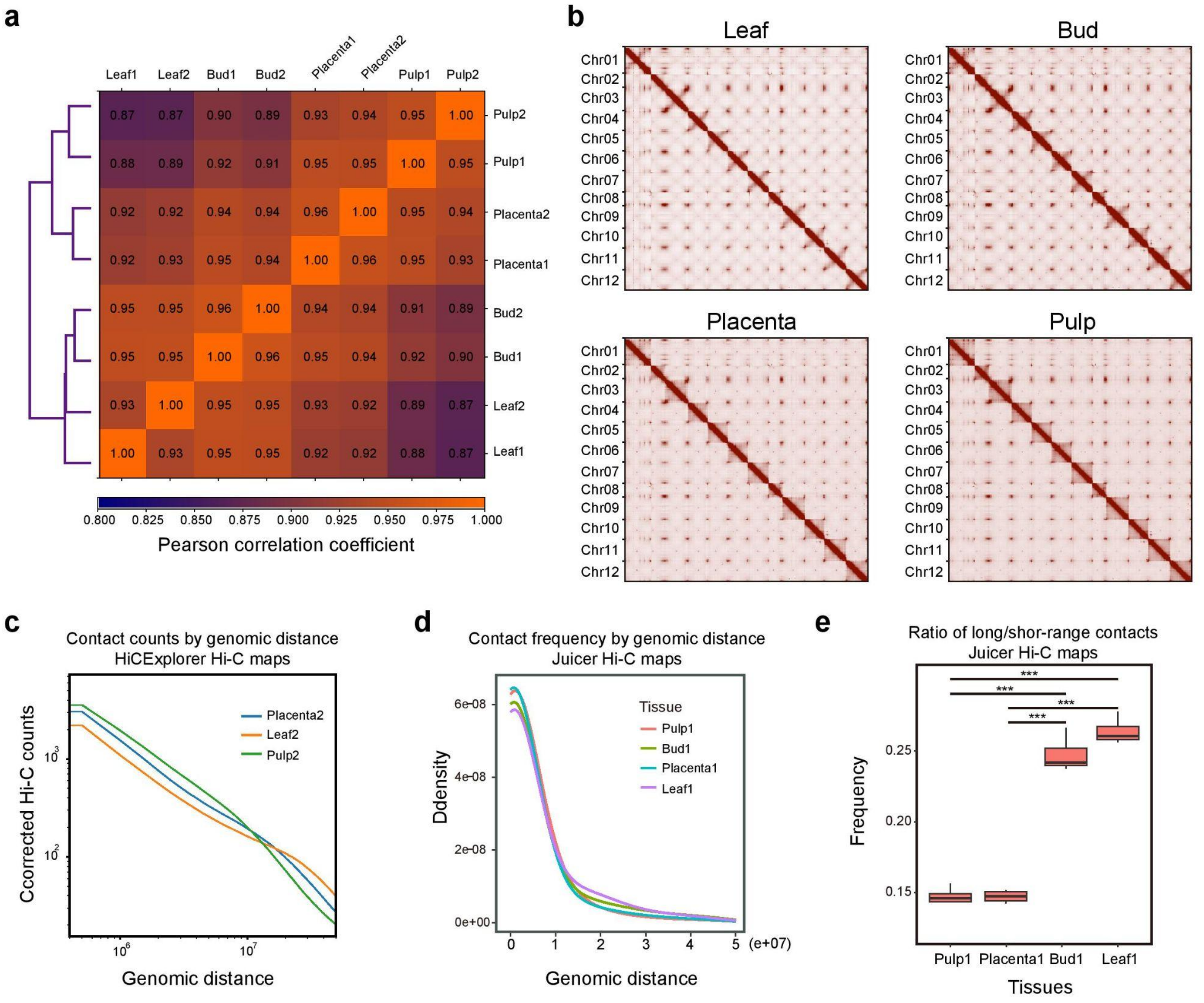


**Extended Data Fig. 5 | Comparison of Hi-C maps among leaf, bud, pulp, and placenta in pepper**. **a,** Pearson correlation analysis of the corrected Hi-C matrices generated by HiCExplorer at 500 kb resolution across samples. **b,** Genome-wide Hi-C heatmaps generated by juicer at 100-kb resolution across four tissues (supplement to Fig. 1a). **c,** The genomic distance *vs.* contact counts plot using Hi-C matrices (HiCExplorer)at 500kb resolution. Samples (leaf, pulp, and placenta) in the second batch were shown (supplement to Fig. 1c). **d,** The genomic distance *vs.* contact counts plot using Hi-C matrices at 500kb resolution generated by juicer pipeline. **e,** The ratio of long-range (>20 Mb) versus short-range contacts for Hi-C matrices generated by juicer was calculated for each chromosome (supplement to Fig. 1d). *** indicates $p < 0.0001$, which were determined by Wilcoxon matched pairs signed rank test.

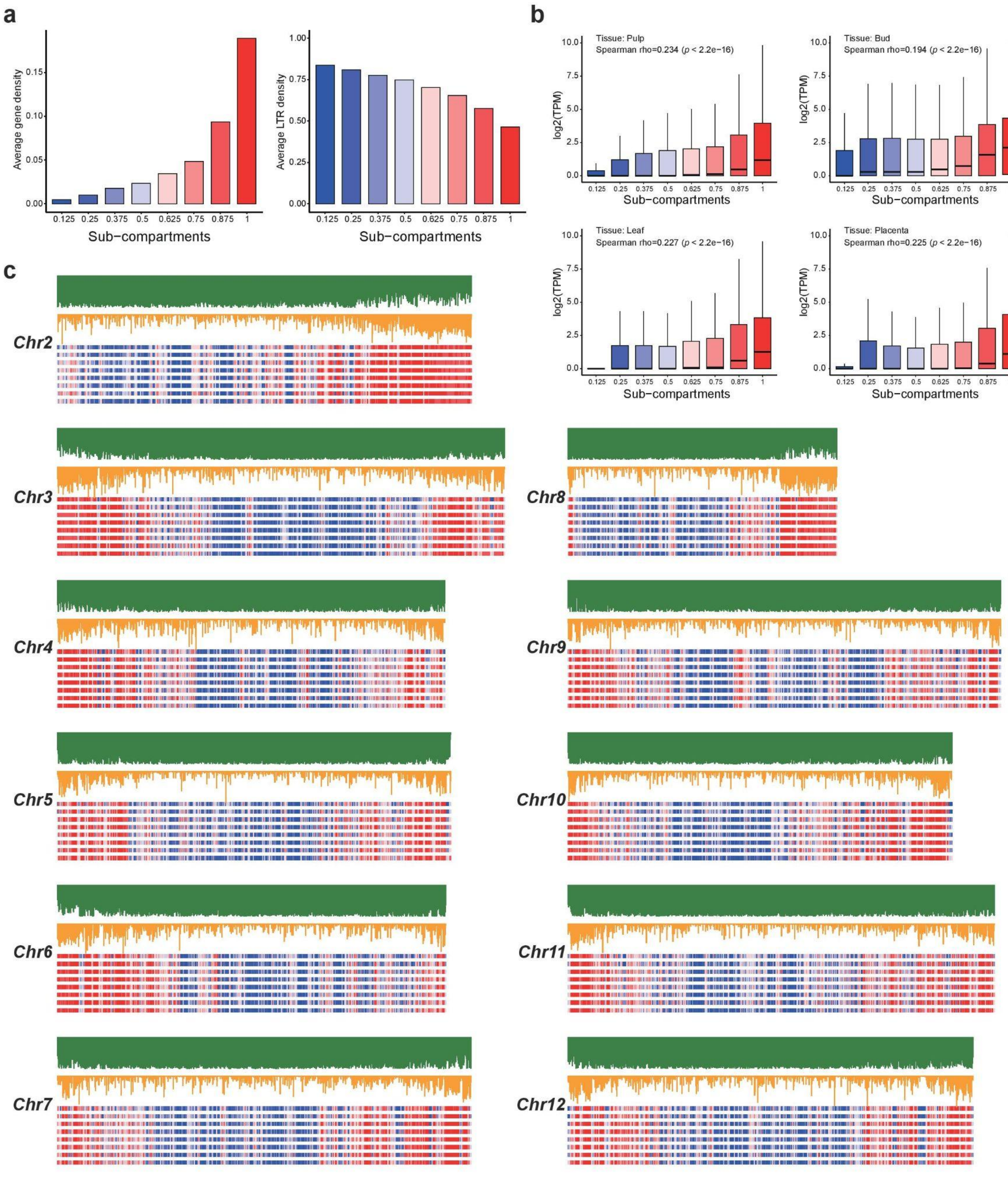
a
Average gene density
0.15
0.10
0.05
0.00
0.125
0.25
0.375
0.5
0.625
0.75
0.875
1
Sub-compartments
Average LTR density
1.00
0.75
0.50
0.25
0.00
b
Tissue: Pulp
Spearman rho=0.234 (p < 2.2e-16)
Tissue: Bud
Spearman rho=0.194 (p < 2.2e-16)
Tissue: Leaf
Spearman rho=0.227 (p < 2.2e-16)
Tissue: Placenta
Spearman rho=0.225 (p < 2.2e-16)
log2(TPM)
10.0
7.5
5.0
2.5
0.0
c
Chr2
Chr3
Chr4
Chr5
Chr6
Chr7
Chr8
Chr9
Chr10
Chr11
Chr12

**Extended Data Fig. 6 | Subcompartments are highly correlated with transcriptional landscape in the pepper genome.** **a**, Calder-inferred subcompartment ranks are positively correlated with gene density (left) but negatively correlated with LTR-RTs density (right). **b**, Subcompartment ranks are positively correlated with transcription levels ($p$-value $<$ 2.2e-16). **c**, Subcompartments are correlated with genome-wide distribution of genes versus retrotransposons (supplement to Fig. 2f).

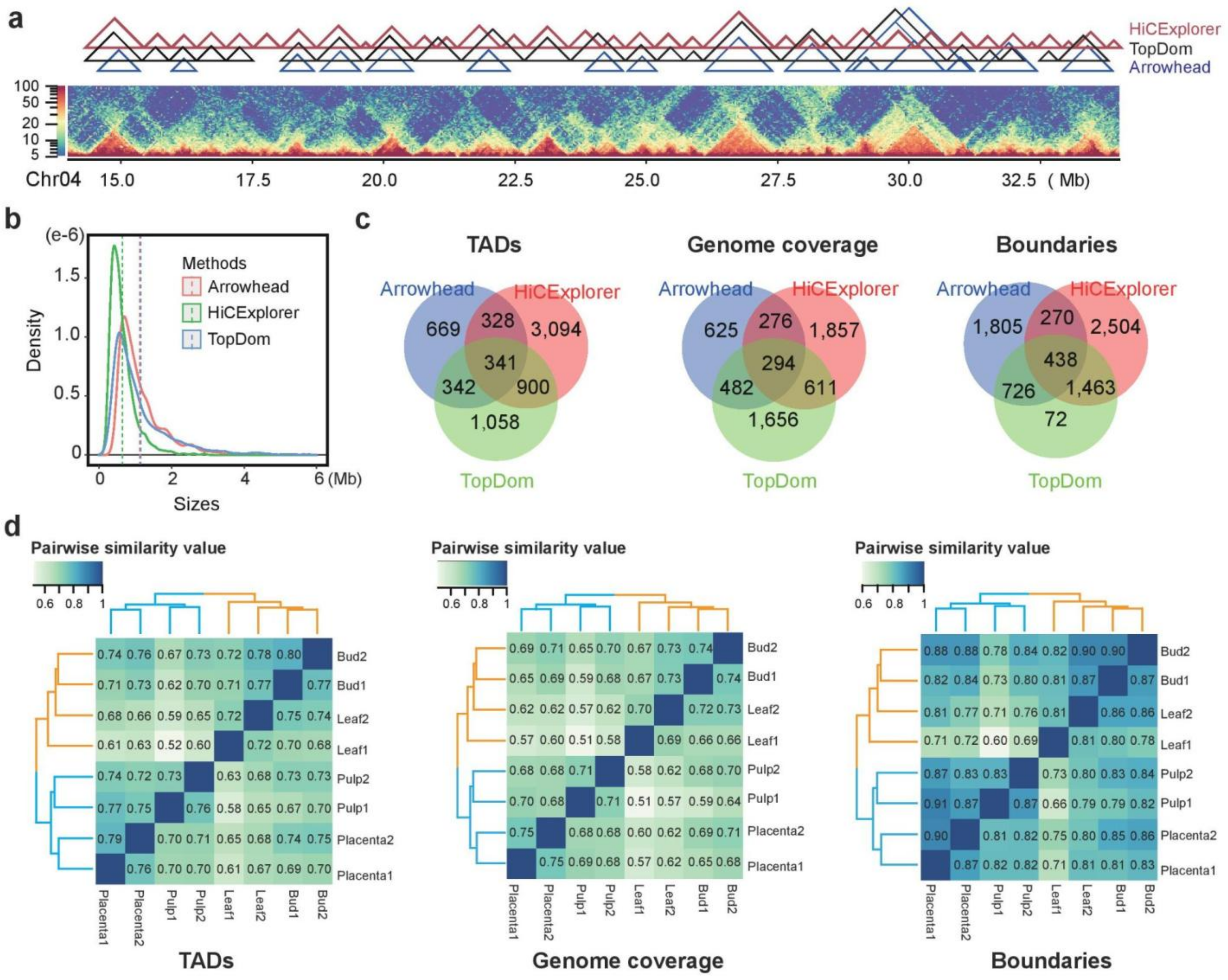


**Extended Data Fig. 7 | Topologically associating domains annotated using different programs, and similarity of TAD structures across tissues. a,** Example of TAD annotation for a 20-Mb region on chromosome 4. TADs were annotated by HiCExplorer, TopDom, and Arrowhead using leaf Hi-C map at 40-kb resolution. **b,** The size distribution of TADs identified by different programs. The mean values were indicated by dashed lines. **c,** Overlap of TADs annotated by different programs, measured in TADs (number), genome coverage, and TAD boundaries. **d,** Hierarchical clustering of samples based on their similarity of TADs, genome coverage, and TAD boundaries. For analyses in **a,b,c**, TAD were annotated from the leaf Hi-C map at 40-kb resolution. For the analysis in **d**, we took TADs from TopDom based on *BNBC* corrected Hi-C maps.

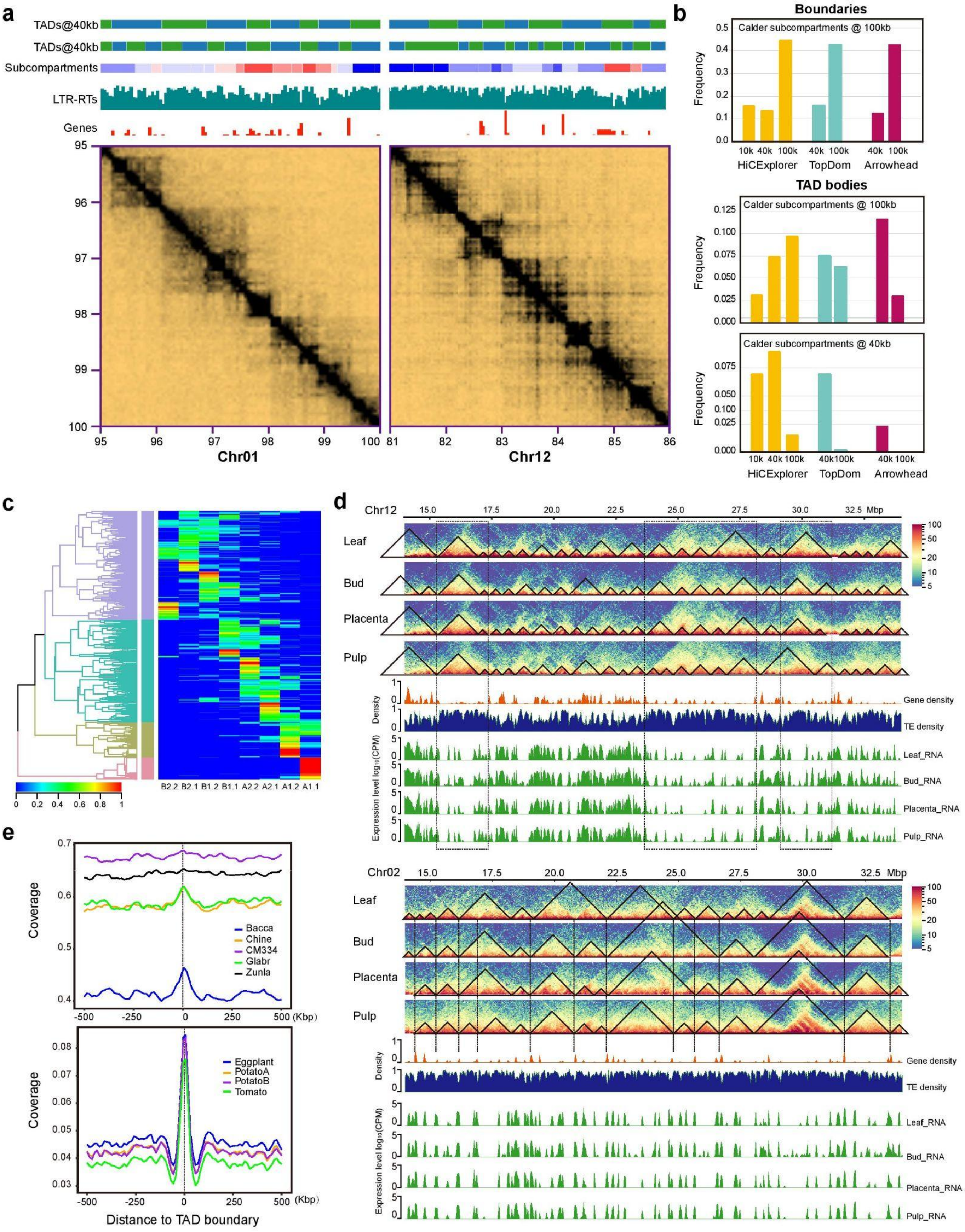

a
TADs@40kb
TADs@40kb
Subcompartments
LTR-RTs
Genes
Chr01
Chr12
b
Boundaries
Calder subcompartments @ 100kb
Frequency
10k 40k 100k
HiCExplorer
TopDom
Arrowhead
TAD bodies
Calder subcompartments @ 100kb
Calder subcompartments @ 40kb
c
B2.2 B2.1 B1.2 B1.1 A2.2 A2.1 A1.2 A1.1
d
Chr12
Mbp
Leaf
Bud
Placenta
Pulp
Density
Gene density
TE density
Expression level log10(CPM)
Leaf_RNA
Bud_RNA
Placenta_RNA
Pulp_RNA
Chr02
e
Coverage
Bacca
Chine
CM334
Glabr
Zunla
(Kbp)
Eggplant
PotatoA
PotatoB
Tomato
Distance to TAD boundary

**Extended Data Fig. 8 | Characterization and classification of topologically associating domains in pepper. a,** Examples of annotation of subcompartments and TADs for two 5-Mb regions on chromosome 1 and 12 (supplement to Fig. 4a). **b,** Assessment of the extent of overlap between TADs and compartmenets/subcompartments, as well as boundaries between them. For boundaries, we compared *Calder*-inferred subcompartment at 100-kb resolution to TAD annotated by HiCExplorer at 10-kb, 40-kb, and 100-kb resolution, and both TopDom and Arrowhead at 40-kb and 100-kb resolution. For TAD bodies, we compared *Calder*-inferred subcompartment at both 40-kb and 100-kb resolution to TAD annotated by HiCExplorer at 10-kb, 40-kb, and 100-kb resolution, and both TopDom and Arrowhead at 40-kb and 100-kb resolution. **c,** Classification of TADs (for those annotated by Arrowhead using Hi-C map at 40-kb resolution) based on the enrichment of *Calder*-inferred subcompartments (supplement to Fig. 4b). **d,** Example of TAD bodies repleting with retrotransposons (above) for a genomic region on chromosome 12, and example of TAD boundaries enriched for genes (below) for a genomic region on chromosome 2. **e,** TAD boundaries are enriched for evolutionary sequence conservation (supplement to Fig. 6b). TAD annotations by HiCExplorer with Hi-C maps at 40-kb resolution were used for analyses in **d** and **e**.

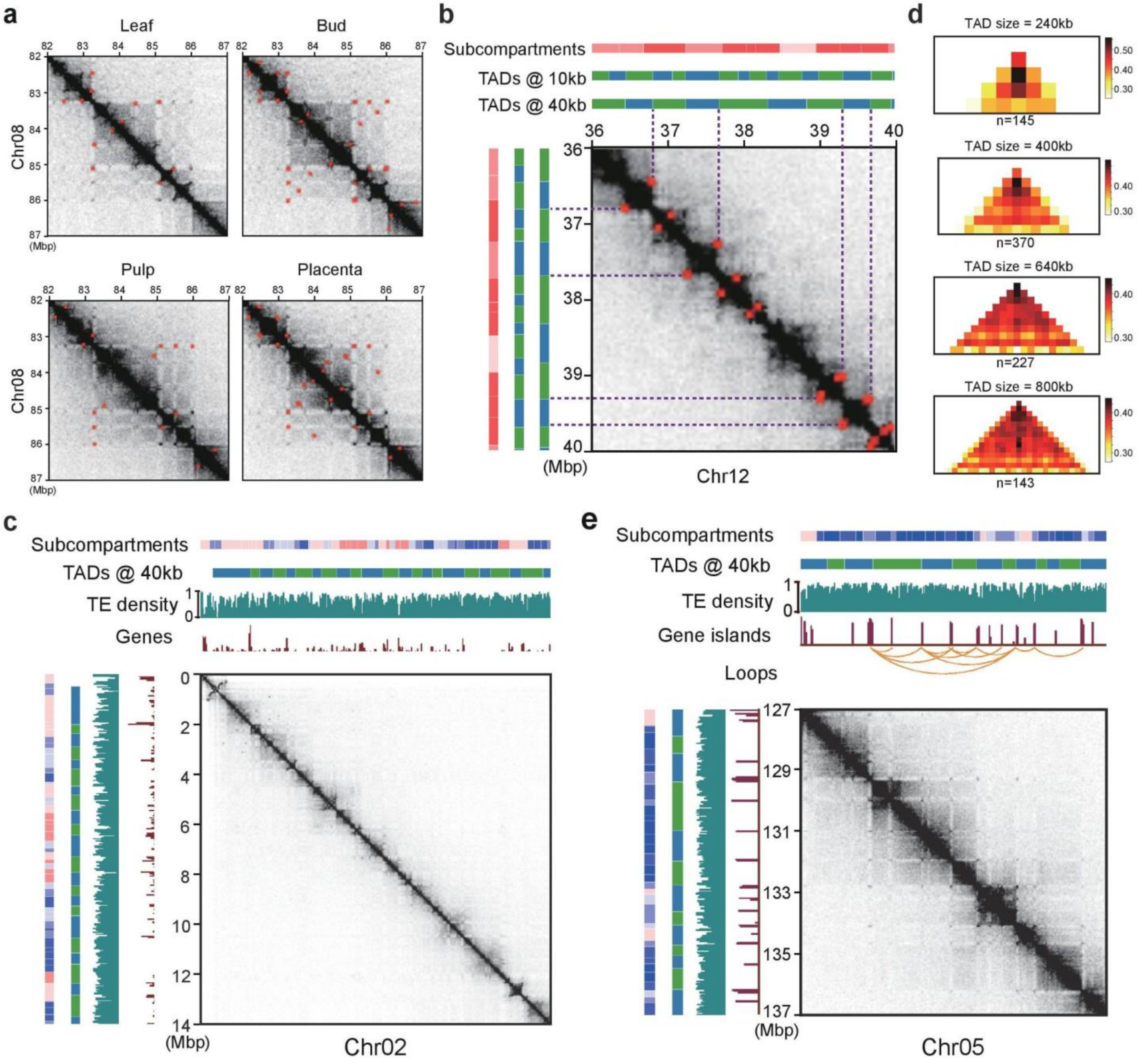


**Extended Data Fig. 9 | Chromatin loops in the pepper genome. a,** Example of loop annotation for a 5-Mb region on chromosome 8 across four tissues. Loops (indicated in red dots) detected in one tissue were missing in another, likely because they were present but below the threshold of detection. Loops were identified by hicDetectLoops. **b,** Example showing a genomic region (Chr12: 36,000,000 - 40,000,000) where chromatin loops demarcate TADs. Subcompartments and TADs identified at both 10-kb and 40-kb resolution for this region were shown above and right (supplement to Fig. 5c). Loops were shown as red dots in the Hi-C contact maps (leaf 40 kb resolution). Dashed purple lines indicate the coincidence of TAD boundaries and loop anchors. **c,** Enhanced contact frequency between the two corners of TADs. TAD sizes are shown on top. Number of TADs for each size is shown below. TADs are identified in the leaf Hi-C map at 40-kb resolution (supplement to Fig. 5d). **d,** Example of Hi-C map showing TADs are demarcated by loops in a gene-rich region on chromosome 2. **e,** Example of gene-to-gene loops for a 10-kb genomic region on chromosome 5 (supplement to Fig. 5e).

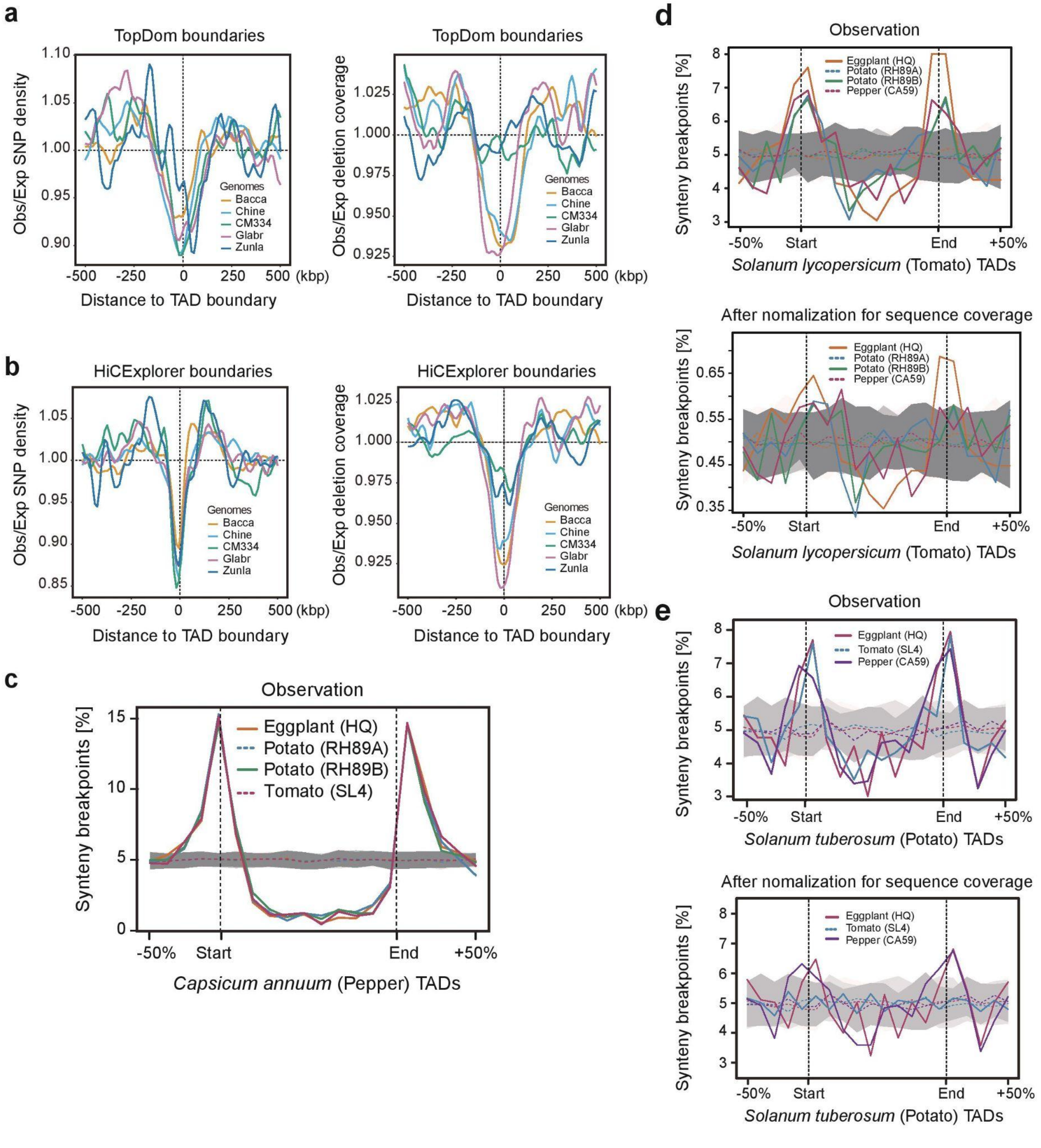


**Extended Data Fig. 10 | Synteny breaks among genomes of solanaceous species are enriched at TAD**

**boundaries, despite evolutionary conservation**. **a,b** The observed (Obs) distribution of SNPs and deletions (coverage) near TAD boundaries relative to the expectation (Exp), based on the genomic background. SNPs and deletions were identified in five closely related genomes (see Fig. 6a) relative to CA59. TADs were annotated by TopDom (**a**) and HiCExplorer (**b**) using leaf Hi-C data at 40-kb resolution (supplement to Fig. 6c). The expected genomic background was calculated as the mean value of all bined windows within 500 kb downstream and upstream of TAD boundaries. **c,** TAD boundaries (observed) of pepper are enriched for evolutionary synteny breaks identified from distantly related solanaceous species (supplement to Fig. 6f). Dotted lines in gray show randomly simulated synteny breaks (n=100). **d,** TAD boundaries of tomato (*Solanum lycopersicum*) are enriched for evolutionary synteny breaks identified from the other three distantly related solanaceous species (eggplant, tomato, and pepper). Top shows the observed values, while bottom shows the normalized values for evolutionary sequence coverage. TADs were annotated by HiCExplorer using Hi-C map at 40 kb resolution. **e,** Similar analyses as **d** when using potato (*Solanum tuberosum*) as the reference.

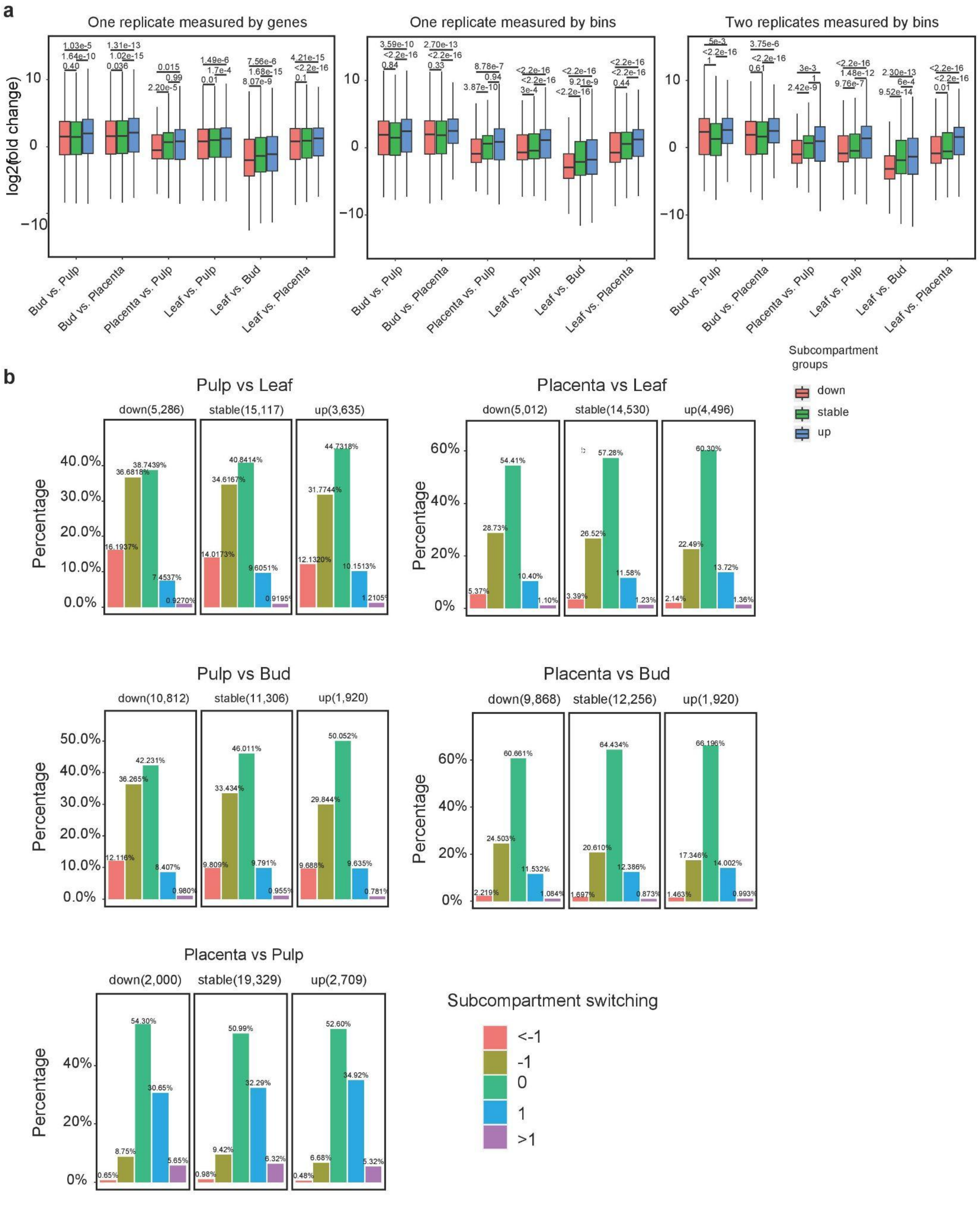
a
One replicate measured by genes
One replicate measured by bins
Two replicates measured by bins
log2(fold change)
Bud vs. Pulp
Bud vs. Placenta
Placenta vs. Pulp
Leaf vs. Pulp
Leaf vs. Bud
Leaf vs. Placenta
Subcompartment groups
down
stable
up
b
Pulp vs Leaf
down(5,286)
stable(15,117)
up(3,635)
Placenta vs Leaf
down(5,012)
stable(14,530)
up(4,496)
Pulp vs Bud
down(10,812)
stable(11,306)
up(1,920)
Placenta vs Bud
down(9,868)
stable(12,256)
up(1,920)
Placenta vs Pulp
down(2,000)
stable(19,329)
up(2,709)
Percentage
Subcompartment switching
<-1
-1
0
1
>1

**Extended Data Fig. 11 | The relationship between subcompartment switching and change in gene expression.** **a,** Genomic regions (i.e. 40-kb bins) switching from A to B compartments or from higher subcompartments to lower subcompartments (e.g. from A1.1 to A1.2) show a trend of decreasing expression, and conversely, switching from B to A compartment or from lower subcompartments to higher subcompartments show a trend of increasing expression . Pairwise comparisons of subcompartment shifts for expression profiles across leaf, bud, pulp and placenta are shown(supplement to Fig. 7a). Analyses were conducted in two ways: 1) only considered one replicate (i.e. a subcompartment switching event only needed to be supported in the first replicate), and 2) two replicates (i.e. a subcompartment switching event needed to be supported by both replicates). The express level was measured in genes or 40-kb bins. **b,** 40-kb bins with decreased expression were slightly enriched for cases of subcompartment switching from a higher rank to lower ranks, while those with increased expression were slightly enriched for cases of subcompartment switching from lower ranks to higher ranks (supplement to Fig. 7b). All reported *P* values are determined by Wilcoxon rank-sum test.

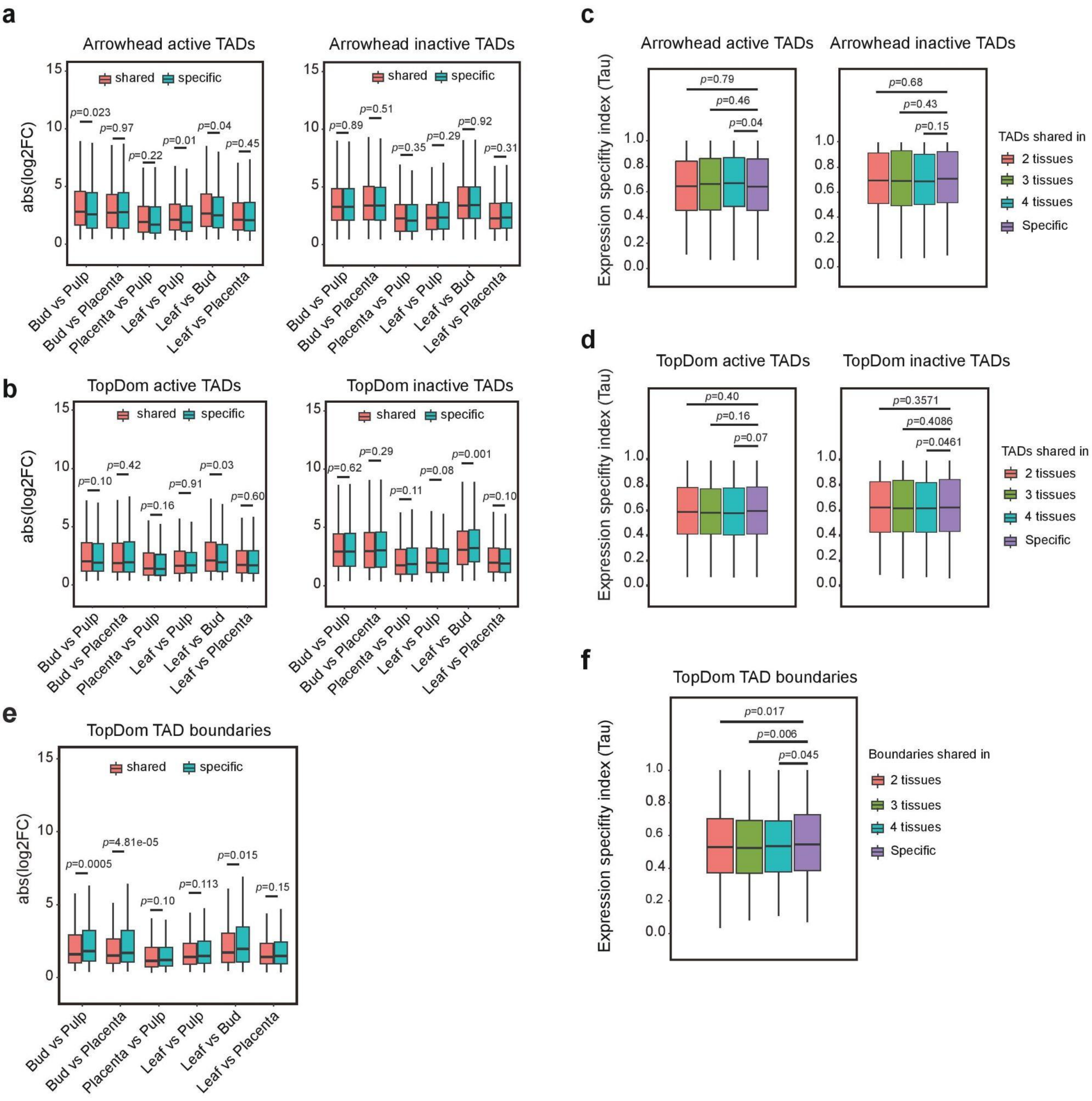
a
Arrowhead active TADs
shared
specific
p=0.023
p=0.97
p=0.22
p=0.01
p=0.04
p=0.45
abs(log2FC)
Bud vs Pulp
Bud vs Placenta
Placenta vs Pulp
Leaf vs Pulp
Leaf vs Bud
Leaf vs Placenta
Arrowhead inactive TADs
p=0.89
p=0.51
p=0.35
p=0.29
p=0.92
p=0.31
b
TopDom active TADs
p=0.10
p=0.42
p=0.16
p=0.91
p=0.03
p=0.60
TopDom inactive TADs
p=0.62
p=0.29
p=0.11
p=0.08
p=0.001
p=0.10
e
TopDom TAD boundaries
p=0.0005
p=4.81e-05
p=0.10
p=0.113
p=0.015
p=0.15
c
Arrowhead active TADs
Arrowhead inactive TADs
Expression specifity index (Tau)
p=0.79
p=0.46
p=0.04
p=0.68
p=0.43
p=0.15
TADs shared in
2 tissues
3 tissues
4 tissues
Specific
d
TopDom active TADs
TopDom inactive TADs
p=0.40
p=0.16
p=0.07
p=0.3571
p=0.4086
p=0.0461
f
TopDom TAD boundaries
p=0.017
p=0.006
p=0.045
Boundaries shared in

**Extended Data Fig. 12 | Conservation of TAD boundaries is associated with transcription stability across tissues but not for TAD bodies**. **a,** Boxplot (all four quartiles shown via lower whisker, lower half of box, upper half of box, and upper whisker;lines indicate median; outliers not shown) showing the *tau* value calculated based on 40 kb bins that fall into TADs shared in 2, 3, and 4 tissues, respectively, compared to tissue-specific TADs. TADs annotated by Arrowhead (leaf Hi-C map at 40-kb resolution) were used in the analysis. TADs are subdivided into active and inactive groups. See **c** for results derived from TADs identified using TopDom. **b,** Boxplot showing the absolute fold change (abs($log_2$FC)) in transcription level (measured in 40-kb bin) for genomic regions that fall into TADs that are shared between tissues compared to genomic regions that fall into TADs that are tissue-specific. See **d** for results derived from TADs identified using TopDom. **e,** 40-kb bins overlapping with shared TAD boundaries (TopDom) across tissues exhibit a significantly lower expression specificity index Tau value compared to those overlapped with tissue-specific TAD boundaries (supplement to Fig. 7d). **f,** 40-kb bins overlapping with TAD boundaries (TopDom) that were conserved between tissues exhibit a relatively smaller absolute change fold in expression level than those overlapping with tissue specific TAD boundaries (supplement to Fig. 7c). Pairwise comparison across four tissues was conducted. All P-values reported were determined by Wilcoxon rank-sum test.
